# Accuracy of haplotype estimation and whole genome imputation affects complex trait analyses in complex biobanks

**DOI:** 10.1101/2022.06.27.497703

**Authors:** Vivek Appadurai, Jonas Grauholm, Morten Krebs, Anders Rosengren, Alfonso Buil, Andrés Ingason, Ole Mors, Anders D. Børglum, David M. Hougaard, Merete Nordentoft, Preben B. Mortensen, Olivier Delaneau, Thomas Werge, Andrew J. Schork

## Abstract

Sample recruitment for research consortia, hospitals, biobanks, and personal genomics companies span years, necessitating genotyping in batches, using different technologies. As marker content on genotyping arrays varies systematically, integrating such datasets is non-trivial and its impact on haplotype estimation (phasing) and whole genome imputation, necessary steps for complex trait analysis, remains under-evaluated. Using the iPSYCH consortium dataset, comprising 130,438 individuals, genotyped in two stages, on different arrays, we evaluated phasing and imputation performance across multiple phasing methods and data integration protocols. While phasing accuracy varied both by choice of method and data integration protocol, imputation accuracy varied mostly between data integration protocols. We demonstrate an attenuation in imputation accuracy within samples of non-European origin, highlighting challenges to studying complex traits in diverse populations. Finally, imputation errors can modestly bias association tests and reduce predictive utility of polygenic scores. This is the largest, most comprehensive comparison of data integration approaches in the context of a large psychiatric biobank.

## Introduction

A recent appreciation for the polygenic nature of complex traits, with several small-effect risk loci scattered throughout the genome has revealed that genome wide association studies (GWAS)^1, 2^ require hundreds of thousands of participants to identify trait-associated loci. Due to their cost-effective nature, genotyping arrays, that ascertain between 200,000 to 2 million single nucleotide polymorphisms (SNPs) in the human genome, have become the preferred technology for generating genetic data at such sample sizes. A key component of these studies is reference-based whole genome imputation (imputation), which expands the number of markers that can be studied^3^, in a two-step process. First, a collection of genotyped SNPs are organized into haplotype scaffolds (phased), relying on co-inheritance patterns of SNPs (i.e., linkage disequilibrium, LD). Known, untyped variants are then probabilistically imputed by matching these sparse scaffolds to more dense haplotypes from whole genome sequenced (WGS) reference individuals^4^. This process results in a much larger pool of variants, thereby increasing GWAS power^5^. Importantly, it helps build a common set of SNPs for meta-analysis across cohorts genotyped on different arrays^6^, and ensures sufficient overlap of SNPs between reference and target datasets for polygenic scoring (PGS)^7^. Various computational methods and reference datasets have been designed for this purpose. Research cohorts beginning with different marker sets, in diverse batches are often combined, even within a single population study.

State of the art phasing methods, such as BEAGLE5^8^, SHAPEIT4^9^ and EAGLE2^10^ use hidden Markov model approaches built on the Li and Stephens model^11^. This model assumes that an individual’s genome can be constructed as a mosaic of segments from haplotypes observed in the reference data or the study population, while accounting for additional factors such as recombination and *de novo* mutation rates. Current phasing methods differ in their computational approximations and data structures used for selecting the most informative haplotypes. Each phasing method further accepts user-defined parameters to choose the number of informative haplotypes, with a trade-off between accuracy, run times, and memory usage. While phasing methods have been improved over the years to scale computationally with large datasets such as the UK biobank^12^, benchmarking is often performed in subsets of the 1000 genomes project^13^, UK biobank, genome in a bottle dataset^14^, or the GERA cohort^15^. To the best of our knowledge, the robustness of these methods has not been tested on input datasets with varying SNP density, target sample sizes, and missingness that can arise when integrating data generated on different genotyping platforms. It is important to empirically characterize the accuracy of phasing and imputation in such scenarios so that researchers can make informed choices when designing bioinformatics workflows to construct next generation biobanks.

The predominant approach used by research consortia for analyzing samples genotyped on multiple arrays has been to phase and impute them separately, prior to meta-analyzing the results for GWAS^16, 17^. However, the accuracy of phasing has been demonstrated to increase with increased sample sizes of reference *and* target datasets^18^. Moreover, for samples generated from recent population-scale biobanks (e.g., UKbiobank^12^, iPSYCH^19^), the number of study individuals is often much greater than the largest available haplotype reference. Haplotype sharing among study individuals and geographical variation in haplotype frequency imply these study haplotypes are as informative, if not more than published references for phasing^20^. Hence, there is intuitive reasoning to pool together as many samples as possible for phasing. In the UK Biobank study, where 500,000 participants were genotyped in 33 batches using two genotyping arrays, it was possible to phase and impute the entire study population together, leveraging the unprecedented sample size because the arrays used, the UK Biobank Axiom array and the UK BiLEVE array, were closely matched (95% marker overlap). However, challenges arise in scenarios where genotyping involves different arrays with low marker overlap and there is currently insufficient guiding research.

Earlier studies on integrating cohorts genotyped on different arrays were on a much smaller scale, used earlier generations of methods, and focused on less diverse cohorts. Sinnott et. al (2012)^21^ compared imputed allele frequencies in two groups of healthy European ancestry controls, genotyped on different arrays with only ∼30% overlap. They observed a substantial type-I error rate, even at genome-wide significance, due to associations with the genotyping array. Retaining only the set of SNPs imputed at the highest quality reduced, but did not eliminate, these errors. Uh et. al (2012)^22^ combined two data sets imputed from arrays with 60,000 overlapping markers into a union data set with high levels of missingness. GWAS across all good quality imputed markers showed an inflation in test statistics that was higher than when restricting to the markers genotyped on both arrays or only including subjects genotyped on one array. The inflation was reduced when an extreme quality control was applied (r^2^ quality metric > 0.98). Johnson et. al (2013)^23^ compared two approaches for integrating cases and controls genotyped on different arrays. They observed that imputing from the union of SNPs across arrays led to 0.2% of SNPs showing associations to genotyping arrays, while imputing from the intersection led to lower imputation accuracy, albeit without the same bias. These previous studies highlight challenges associated with integrating genotype data, including the important notion of a potential accuracy/bias trade-off, but do not provide a consensus path forward.

Pimental et. al^24^ studied the biases introduced by imputation in the context of estimating direct genomic values (DGV) in livestock, analogous to PGS in human genetics. They observed a bias in imputed genotypes towards the more frequent (major) allele in the reference panel that caused estimated DGVs to be shrunk towards the sample mean. This bias was more evident in traits with high heritability and when DGVs were estimated using imputation from less dense haplotypes. More recently, Chen et. al^25^ studied the impact of different combinations of phasing and imputation methods on PGS and demonstrated that while PGS differ at an individual level, when computed using imputed genotypes rather than gold standard WGS, the variation at cohort level is low, resulting in a less than 5 percentile change in individual PGS rank within the cohort. The impact of imputation on PGS in context of data integration across cohorts has otherwise remained underexplored and given the attention PGS have recently received^26–29^, exploring these concepts in modern, population-scale, human complex trait genetics applications is critical.

This study uses the Lundbeck foundation initiative for integrative psychiatric research (iPSYCH) case-cohort dataset with an initial 81,330 subjects genotyped on the Infinium PsychChip v1.0 (Illumina, San Diego, CA USA) and an additional 49,108 subjects genotyped on the Illumina Global Screening Array v2.0 (Illumina San Diego, CA USA) to evaluate four realistic protocols for data integration. We compare the phasing accuracy using SHAPEIT4.1.2, EAGLE2.4.1, BEAGLE5, and a consensus approach in truth sets derived from 124 parent-offspring trios that were genotyped on both arrays. To compare the resulting imputation quality, we randomly masked 10,000 SNPs prior to phasing and included 10 WGS samples from the Personal Genomes Project - UK cohort^30^, down sampled to the SNPs in each cohort. Imputed genotypes were then compared to these truth sets to assess the loss of information in imputed data. It is known that current haplotype references are skewed towards individuals of European ancestry, hence we assessed quality of phasing and imputation in non-European and admixed individuals. Finally, using a simulated quantitative trait, we explore the impact of phasing and imputations across data integration scenarios on GWAS and PGS.

## Materials and methods

### Data

iPSYCH2012 is a case-cohort design nested within 1,472,762 individuals born in Denmark between 01-05-1981 and 31-12-2005, with a known mother, alive and residing in Denmark at the end of the first year after birth. Out of 86,189 individuals chosen for genotyping, 57,377 are cases with one or more mental disorders among schizophrenia, autism, attention-deficit/hyperactivity disorder (ADHD) and affective disorder. The cohort is a random sample of 30,000 individuals representative of the national population of Denmark born during the same time period. Genotyping was performed at The Broad Institute, Boston MA, USA with the Infinium PsychChip v1.0 (Illumina, San Diego CA, USA), using DNA extracted from dried blood spots, obtained from the Danish neonatal screening biobank^31^. Further details on the ascertainment and data generation process of iPSYCH2012 has previously been described^19^. iPSYCH2015i is an extension of iPSYCH2012, nested within 1,717,316 individuals born in Denmark between 01-05-1981 and 31-12-2008, satisfying the same criteria, encompassing 33,345 cases and 15,756 cohort individuals, genotyped on the Illumina Global Screening Array v2.0 (Illumina, San Diego CA, USA) at Statens Serum Institut, Copenhagen Denmark.

The trio dataset contains 128 parent-offspring trios where the offspring were ascertained for diagnoses of autism or ADHD with both parents born in Denmark, on or after 01-05-1981. Samples were genotyped using both the Infinium PsychChip v1.0 and the Illumina Global Screening Array v2.0. Information on psychiatric diagnoses were obtained from the Danish national psychiatric central register^32, 33^, demographic information including age, gender and parental birth place were obtained from the Danish civil registration system^34, 35^.

The Personal Genomes Project - UK (PGP-UK) is an open source initiative aimed at facilitating access to multi-omics datasets for the purpose of gaining insights into biological and medical processes^30^ and contains 1,100 citizens or permanent residents of the United Kingdom who provided consent after passing a test aimed at educating them on the risks of sharing personal genetic data. DNA was extracted from blood and whole genome sequenced using Illumina HiSeq X at an average depth of 15x. The resulting BAM files were deposited to the European Nucleotide Archive (Study identifier: PRJEB17529).

### Ethical Permissions

Research using iPSYCH and the trio data has been approved by the Danish scientific ethics committee, Danish health authority and the Danish neonatal screening biobank committee. PGP-UK has been approved by the University College London scientific ethics committee. All analyses were performed on a secure server within the Danish national life science supercomputing cluster (https://computerome.dtu.dk/) and the Aarhus Genome Data Center (https://genome.au.dk/).

### Genotype Quality Control (QC)

Genotype data from iPSYCH2012, iPSYCH2015i, trios and PGP-UK were aligned to HRC v1.1 using genotype harmonizer version 1.4.20-SNAPSHOT^36^. SNPs not genotyped in all waves/batches within individual iPSYCH cohorts were excluded. Further filtration steps include exclusion of SNPs missing in at least 5% of the study subjects, SNPs showing differential missingness between cases and controls, SNPs failing tests of Hardy Weinberg equilibrium in controls of a homogenous genetic origin, SNPs significantly associated with a genotyping batch or wave, SNPs with minor allele frequencies less than 0.001. A further 10,000 SNPs were selected at random and masked to serve as a truth set for benchmarking the performance of imputation. Samples were excluded if they had abnormal levels of heterozygosity that could not adequately be explained by admixture, or if missing more than 5% of the SNPs that passed QC. In case of duplicate samples or monozygotic twins, the sample with lower missingness was retained. This resulted in a total of 80,876 individuals genotyped at 251,551 SNPs in iPSYCH2012 and 48,974 individuals genotyped at 450,445 SNPs in iPSYCH2015i passing QC. QC detailed in depth in supplementary S1. PLINK v1.90b3o 64-bit 20 May 2015^37^ was used for QC.

### Pre-phasing Data Integration Protocols

We evaluated four different ways of integrating data as shown in Figure 1.

**Figure 1.**
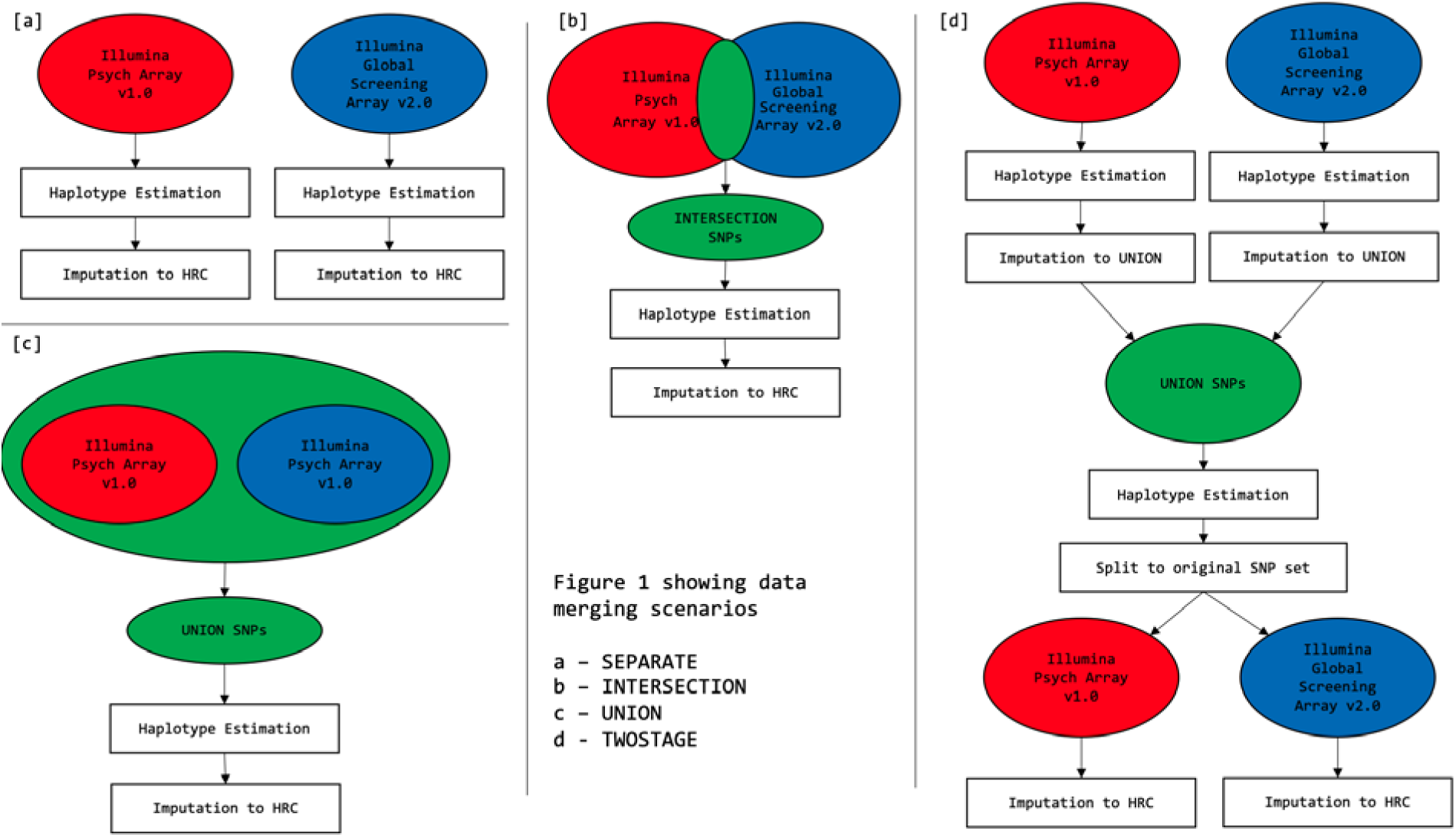
Four pre-phasing data integration protocols. [a] shows the *separate* protocol where the cohorts genotyped on each array are phased and imputed separately. [b] shows the *intersection* protocol where the two cohorts are merged to include only SNPs in common to both genotyping arrays prior to phasing and imputation. [c] shows the *union* protocol where the two cohorts are merged to include SNPs genotyped on either array and the resulting dataset with missingness is phased and imputed. [d] Shows the *twostage* protocol where the haplotypes obtained from the *separate* protocol are initially imputed to the markers in the *union* protocol, prior to a second stage of phasing before the cohorts are split back to the original sets of genotyped SNPs after which imputation to the full reference panel is performed.

#### Separate

In this protocol (Figure 1a), samples from iPSYCH2012 and iPSYCH2015i are phased and imputed separately. 124 trio offspring were added to both cohorts. Ten whole genome sequenced samples from the PGP-UK cohort were down sampled to both the iPSYCH2012 and iPSYCH2015i SNPs that passed QC and merged with both cohorts. This resulted in two cohorts: (1) Cohort2012 (81,022 samples, 251,551 SNPs, 0.1% missingness) which includes iPSYCH2012, trio offspring genotyped on the Infinium PsychChip v1.0 and ten PGP-UK samples, down sampled to the Infinium PsychChip v1.0 variants that pass QC. (2) Cohort2015i (49,120 samples, 450,455 SNPs, 0.31% missingness) which includes the iPSYCH2015i, trio offspring genotyped on the Illumina Global Screening Array v2.0 and ten PGP-UK samples, down sampled to the Illumina global screening array v2.0 variants that pass QC.

#### Intersection

In this protocol (Figure 1b), samples from iPSYCH2012 and iPSYCH2015i were merged at the 116,962 QC’ed SNPs present on both iPSYCH arrays. 62 offspring samples were chosen at random from each of the trio datasets genotyped using both iPSYCH arrays and merged to this dataset along with ten PGP-UK samples, down sampled to the 116,962 common loci. This resulted in the *intersection* (129,886 samples, 116,962 SNPs, 0.17% missingness) cohort.

#### Union

In this protocol (Figure 1c), samples from iPSYCH2012, iPSYCH2015i were merged with missingness to the 596,028 QC’ed SNP loci, genotyped on either iPSYCH array. To this, 62 samples each from the trio dataset genotyped on both arrays were merged, same as in the *intersection*. Five PGP-UK samples, each down sampled to the SNPs present on either genotyping array, were merged resulting in the *union* cohort (129,886 samples, 596,028 SNPs, 44.54% missingness).

#### Two-stage

In this protocol (Figure 1d), eight sets of phased haplotypes from the Cohort2012 and Cohort2015i obtained in the *separate* protocol were initially imputed using BEAGLE5.1 in batches of 10,000 samples to the 596,028 QC’ed SNPs genotyped on either iPSYCH array with HRCv1.1 as the reference. Then the two cohorts were merged, retaining the same 62 trio samples from each cohort as chosen in the *intersection* and *union* approaches along with five PGP-UK samples from each cohort, forming the *twostage* cohort (129,886 samples, 596,028 SNPs, 0% missingness).

All datasets were stored and processed in variant call format (VCF) (http://samtools.github.io/hts-specs/VCFv4.2.pdf) using bcftools^38^.

### Phasing

Cohorts arising from each data integration protocol were phased using three methods and two different parameters, BEAGLE5 (phase-states=280, 560), SHAPEIT4.1.2 (pbwt-depth = 4, 8), EAGLE2.4.1 (Kpbwt = 10000, 20000) with the added aim of benchmarking improvements in accuracy at a higher resolution parameter set at the expense of longer run times and memory requirements. A *consensus* haplotype set was generated by taking the majority haplotype estimate across the three tools at both the default and higher resolution parameters at each locus within each individual using BEAGLE’s *consensusvcf* module (<u>consensusvcf.jar</u>). The HRCv1.1 dataset, consisting of 64,976 haplotypes^39^ was used as the reference panel.

### Imputation

All cohorts were imputed using BEAGLE5.1 with HRCv1.1 as the reference. Due to the cohort sizes, imputations were carried out in batches of 10,000 samples. Imputed dosage (DS) for an individual at a bi-allelic locus is calculated as DS = p(RA) + 2*p(AA) where p(RA) is the genotype probability corresponding to the presence of one alternate allele (A) and one reference allele (R) as per the reference panel and p(AA) corresponding to the genotype probability of the presence of two copies of the alternate allele.

### Phasing accuracy

Phasing accuracy was evaluated by calculating switch error rates (SER) in the trio offspring at the QC’ed heterozygous SNPs common to both iPSYCH arrays. A switch error occurs when there arises an inconsistency between the computationally assigned phase and the phase observed by mendelian transmission with knowledge of parental haplotypes. SER is the number of such switches divided by the total possible switches^40^. The code for SER calculation has previously been used^9^ and available on GitHub (https://github.com/odelaneau/switchError).

### Imputation accuracy

Imputation accuracy within iPSYCH was calculated as the squared Pearson correlation coefficient (r^2^) between true genotypes and imputed dosages within different minor allele frequency (MAF) bins (MAF as measured in HRCv1.1) at each of the 10,000 SNPs masked prior to phasing. Imputation accuracy within PGP-UK was calculated as the r^2^ between true genotypes obtained from multisampling variant calling using *samtools*^38^ and imputed dosages in eight MAF bins at 6,517,513 loci that were genotyped on neither iPSYCH array. The code is available on GITHUB (https://github.com/vaqm2/impute_paper/blob/main/truth_vs_impute_2021_02_24.pl). To evaluate variations in imputation accuracy by ancestral origin, r^2^ was calculated within iPSYCH samples, grouped according to the country of birth of both parents according to the Danish civil register^34, 35^.

### Phenotype simulations

To evaluate the impact of whole genome imputation on polygenic scores, a quantitative trait for 129,850 iPSYCH individuals was simulated using GCTA^41^ version 1.92.1beta6, with a heritability of 0.5 and the 10,000 masked SNPs as causal loci with effect sizes drawn from a standard normal distribution.

### Association Tests

To evaluate the presence of batch artifacts in each protocol we conducted multiple GWAS with iPSYCH cohort membership (iPSYCH2012 vs iPSYCH2015i) as the outcome using the glm module of PLINKv2.00a2LM 64-bit Intel (10 Nov. 2019)^42^. As a baseline, we performed the GWAS using true values of 10,000 masked genotypes as explanatory variables. Subsequently, GWAS were performed comparing allele frequencies from true genotypes in one cohort to imputed dosages in the other and imputed dosages in both, across all four data integration protocols (*separate*, *union*, *intersection*, *twostage*). Tests were restricted to iPSYCH individuals without mental disorders (i.e., a random sample of psychiatric controls), of a homogenous genetic origin based on principal component analysis (Supplementary S1.1) using Eigenstrat^43^, and pruned for relatedness beyond the third degree using kinship coefficients estimated by KING^44^. The overall inflation of test statistics above the null was evaluated using the genomic inflation factor which compares the median of the chi-square test statistic obtained from each GWAS to the expected median of a chi-square distribution with 1 degree of freedom.

### Polygenic Scores

Polygenic scores (*PGS*) for each individual, *j*, were constructed using simulated per-allele effects as follows:

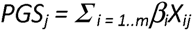

where *m* is the total number of SNPs (10,000 masked SNPs), *β* is simulated effect for SNP *i*, *X_ij_* is the imputed dosage or best guess genotype count of effect alleles for individual j at SNP i. Variance explained by PGS was calculated by fitting two linear models using the function, lm in R. The simulated trait value is the outcome, individual PGS is the sole explanatory variable in one model, while individual PGS, age, gender and first 10 principal components of genetic ancestry are explanatory variables in the second model. Variance in the simulated trait value explained by PGS is the difference between the correlation coefficients (R^2^) between the two models. We restrict the analysis to 67,587 individuals from iPSYCH2012 and 41,069 individuals from iPSYCH2015i with parents and both sets of grandparents born in Denmark and clustering with the CEU (Utah residents with Northern and Western European ancestry) and GBR (British in England and Scotland) populations of the 1000 genomes phase 3 dataset in principal component analysis (Supplementary S1.1).

## Results

### Phasing Accuracy

Phasing accuracy was measured using SER (Methods) with three methods, two parameter settings each, and a consensus set across four data integration protocols (Figure 2a, Supplementary S3, Supplementary Table 9). Our results show that phasing accuracy depends on the data integration protocol, phasing methods and associated parameters, target sample size, genotyped SNP density in the target, rate, and structure of genotype missingness. In general, the *two-stage* protocol, which leverages the largest possible sample size and density of SNPs, with no missingness, shows consistently high accuracy across all phasing methods (SER = 0.17 - 0.55%). The *intersection* protocol, which also leverages the largest sample size, albeit with lowest SNP density, proves the least accurate (SER = 0.38 - 1.04%). The ranking of the protocols was generally consistent across methods, except for the *union*, which achieved the lowest overall SER with BEAGLE5 at parameter value, phase-states=560. The *union* was also the worst performing protocol when taking consensus haplotypes across all three methods (SER = 0.61% at default parameters), suggesting the genotype missingness introduced by this protocol causes systematic phasing errors that are reproduced across tools.

**Figure 2.**
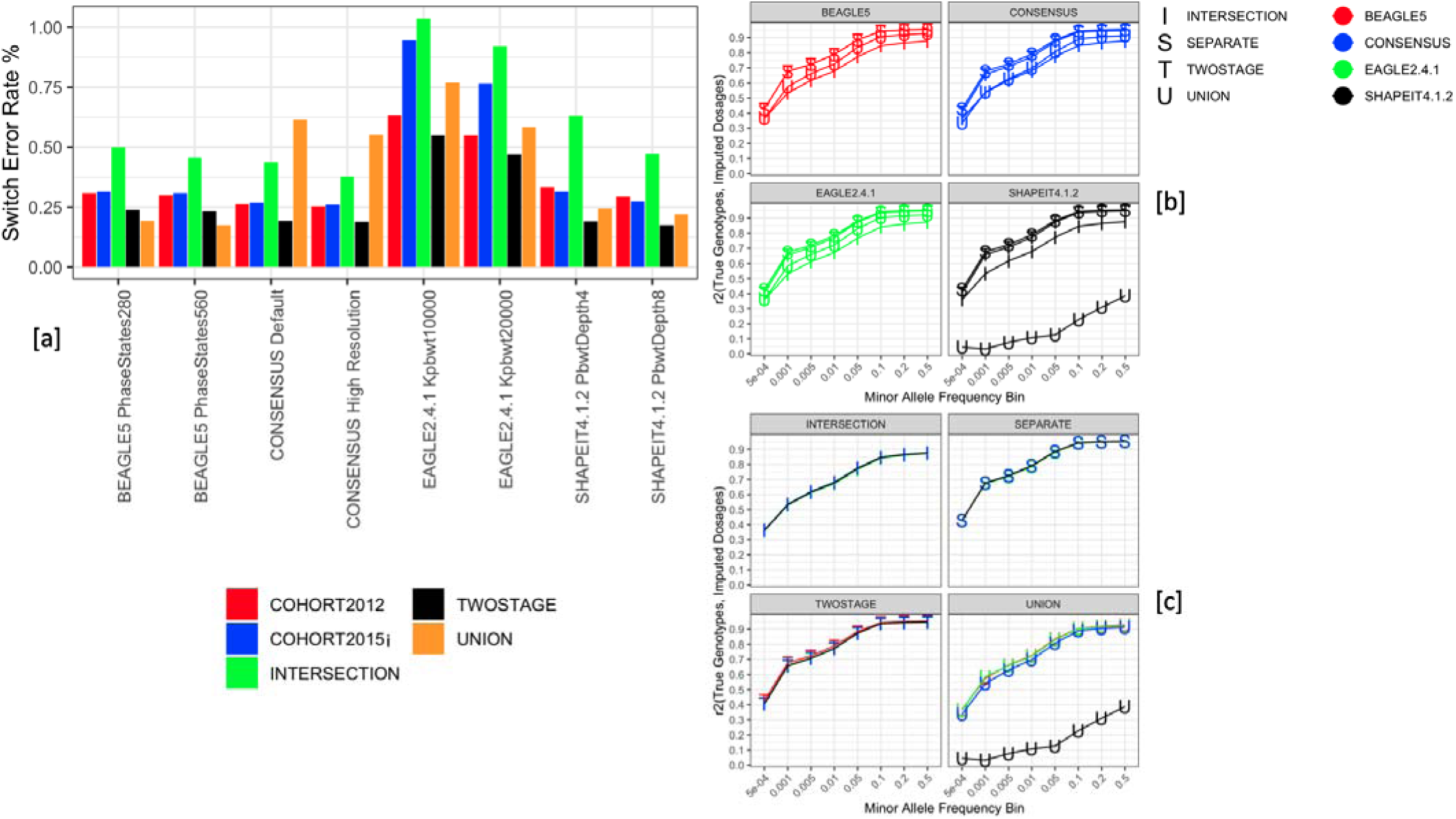
Phasing and imputation accuracy vary across state-of-the-art tools and data integration approaches. [a] Shows the accuracy in switch error rate percentage of phasing across the three tools at two parameter sets each and a *consensus* approach taking the majority haplotype at each locus from the three tools at both parameter sets across all four data integration protocols. Default parameters are SHAPEIT4.1.2 pbwt-depth=4, BEAGLE5 phase-states=280, EAGLE2.4.1 Kpbwt=10000. High Resolution parameters are SHAPEIT4.1.2 pbwt-depth=8, BEAGLE5 phase-states=560, EAGLE2.4.1 Kpbwt=20000. The switch error rates were computed within 124 trio offspring by comparing the computationally assigned phase to the mendelian transmission from known parental genotypes at the heterozygous loci common to both genotyping arrays. [b] Shows the imputation accuracy (r^2^) within each data integration protocol, grouped by choice of phasing tool at different minor allele frequency bins at the 10,000 SNPs common to both genotyping arrays that were masked prior to phasing. [c] shows the accuracy (r2) of imputation from haplotypes estimated using the three different tools and the consensus approach grouped by choice of data integration protocol at different minor allele frequency bins. All imputations were performed using BEAGLE5.1 with the HRCv1.1 as the haplotype reference panel.

In protocols involving little to no genotype missingness (i.e., not *Union*), BEAGLE5 and SHAPEIT4.1.2 show similar accuracy, outperforming EAGLE2.4.1 across integration methods and parameters. The *union* was again a point of departure from the trends, with BEAGLE5 performing better (SER = 0.17%) on the *union* and SHAPEIT4.1.2 performing better on the *twostage* (SER = 0.17%). This implicates genotype missingness for phasing performance, suggesting that BEAGLE5 handles this more robustly than SHAPEIT4.1.2. When considering the *twostage* protocol, which we hypothesized could mitigate initial missing genotypes, SHAPEIT4.1.2 performed like BEAGLE5 on the *union* (and better than on the *twostage*), suggesting, modulo initial missingness, SHAPEIT4.1.2 may have at least as good a phasing algorithm as BEAGLE5.

Comparing the phasing accuracy across chromosomes within each method and data integration protocol reveals that phasing accuracy follows the number of SNPs per centimorgan in the target dataset, with denser chromosomes showing lower SER (Supplementary figure 1). We also observe that EAGLE2.4.1 and BEAGLE5 produce more accurate estimates in the Cohort2012 where the sample size is higher and SNP density is sparser whereas SHAPEIT4.1.2 produces more accurate estimates in the Cohort2015i where the SNP density is higher and target sample size is comparatively smaller. As mentioned above, the worse performance of SHAPEIT4.1.2 and EAGLE2.4.1 on the *union* as opposed to the *twostage* highlight the sensitivity to initial missing genotypes. These results show the necessity for benchmarking the robustness of phasing methods in less-than-ideal conditions, specific to study cohorts, prior to deploying them in such untested scenarios.

### Imputation Accuracy

The accuracy of imputations derived from each set of haplotype scaffolds (i.e., from each tool, parameters and data integration protocol set) are presented in figures 2b, c and Supplementary Table 7. Variability in imputation accuracy stems more from the choice of data integration protocol, rather than the choice of phasing method or parameters. Since all methods process data in variant call format (VCF), this renders the choice of phasing method less relevant if the end goal is to attain the most accurate missing data imputation. The highest imputation accuracy is obtained when the cohorts are phased separately, with the r^2^ between true masked genotypes and imputed dosages varying between 0.43 at rare (MAF < 0.005) and 0.95 at more common (0.2 < MAF <= 0.5) SNPs. This trend is consistent across haplotypes generated by all methods. The added bioinformatics effort aimed at enhancing sample size without missingness with the *twostage* protocol did not yield a higher imputation accuracy than the *separate* protocol. At the minor allele frequency bin, 0.01 < MAF <= 0.05, using haplotypes phased by BEAGLE5, both approaches show identical accuracy (r^2^=0.88) (Supplementary Table 7). The imputation accuracy is degraded when using the *intersection* protocol with an attenuation between 8.4-13.6% at common and 13.9-18.6% at rare SNPs as compared to the *separate* protocol, highlighting the drop in phasing accuracy at low target SNP density carrying over to imputation performance.

Haplotypes estimated by SHAPEIT4.1.2 in the *union* protocol are an outlier and resulting imputations are of noticeably poorer quality compared to haplotypes obtained from other methods. Phasing in the presence of missingness is itself a two-step process, where each phasing method makes a rough imputation of missing data prior to constructing haplotypes. If this data is not overwritten during imputation, the prephasing imputation algorithm implemented by SHAPEIT4.1.2 could be the reason for problems with the *union* protocol. This becomes more credible when considering the imputation accuracy obtained from the *twostage* protocol using SHAPEIT4.1.2, where the attenuation is mitigated. The pattern of results described above is replicated in the PGP-UK samples (Supplementary Figure 2).

A comparison of imputation accuracies between Cohort2012 and Cohort2015i within the *separate* protocol using the PGP-UK samples (Supplementary Figure 2c, Supplementary Table 9) shows higher imputation accuracy in Cohort2015i, imputed from a higher SNP density as compared to Cohort2012 with a larger sample size with a difference as high as 6.7% at E (0.1, 0.05]. This finding is important because it emphasizes a trade-off between sample size and SNP density and, with modern samples, perhaps SNP density should be emphasized. Enhanced parameters that showed higher phasing accuracy do not seem to substantially increase imputation accuracy (Supplementary figures 2a, b). Taken together, these results show that imputation performance suffers when merging cohorts genotyped on different arrays prior to phasing and choice of phasing method is less relevant than data integration protocol.

### Imputation accuracy in non-European and admixed samples

It is known that GWAS results and subsequent PGS constructed from them do not generalize well across populations^45^. This is typically attributed to inaccuracies in the estimation of SNP effect sizes (i.e., per SNP beta) due, e.g., to variable LD across populations^46^. However, if non-European haplotypes are underrepresented in either reference or target data sets, imputed genotypes in these individuals may be of lower quality and errors in the genotypes themselves could be contributing to the generalization problems of GWAS. Imputation accuracy was estimated in non-European and admixed iPSYCH samples, grouped according to the birthplace of the proband’s parents (Figure 3a, b; Supplementary Figure 4, Supplementary Table 8). Individuals born to non-Scandinavian European parents had lower imputation accuracy (7.07-12.58%) than those with both parents born in Denmark. These effects were larger for individuals with both parents born in Asia (11.1-11.2%), Africa (17.37-17.48%), or Middle East (11.2-17.7%). The attenuation in imputation accuracy within admixed individuals is comparatively lower, varying between 4.47-8.56% as compared to individuals with both parents born in Denmark. These results, as expected, suggest that imputation accuracy varies by ancestry and introduces a systematic loss of information in the genotypes of non-Europeans.

**Figure 3.**
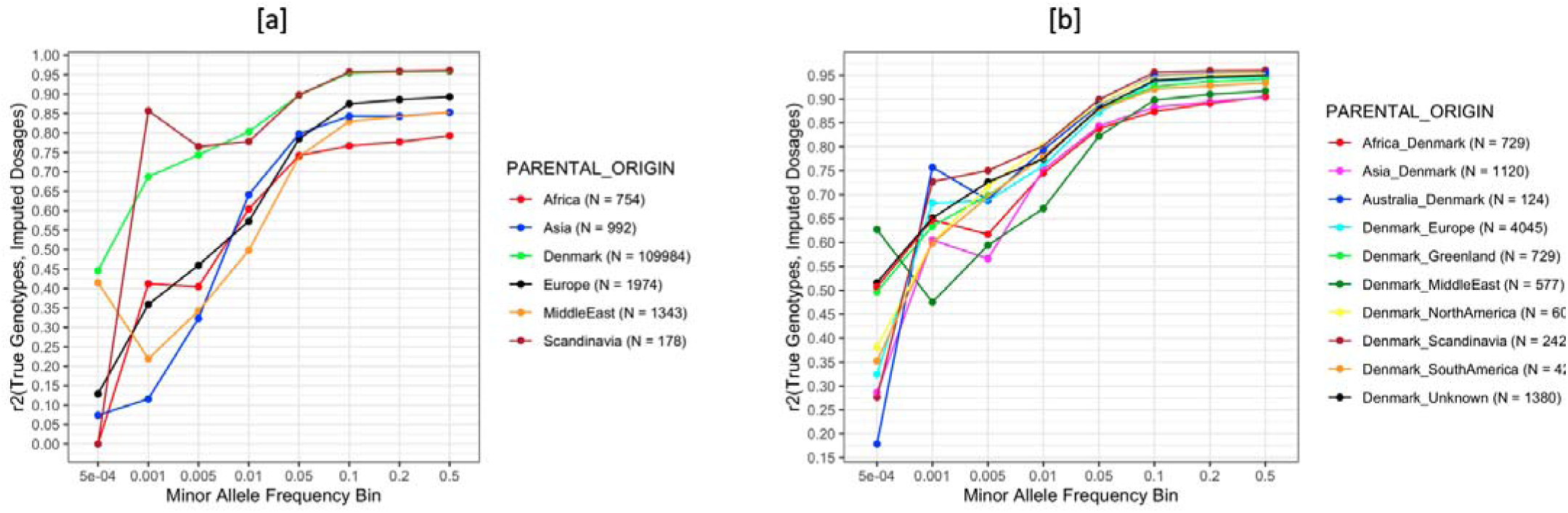
The accuracy of imputation varies extensively by genetic ancestry. [a] shows the imputation accuracy (r^2^) in iPSYCH samples grouped by parental birthplace as ascertained from the Danish civil registers at different minor allele frequency bins within the 10,000 SNPs common to both genotyping arrays, masked prior to phasing. [b] shows the imputation accuracy (r^2^) in admixed samples where at least one parent was born in Denmark. All imputations were performed using the *separate* protocol, haplotype estimation was performed using BEAGLE5 phase-states=560, imputations were performed using BEAGLE5.1 with the HRCv1.1 as the reference.

### Impact on PGS

PGS calculated using true genotypes in iPSYCH2012 and 2015i respectively explained 45.4 and 45.6% of the variance in the simulated continuous trait (Figure 4a, Supplementary S7). An attenuation in variance explained was observed when PGS were instead calculated using imputed dosages, which was lowest when using the *separate* protocol for iPSYCH2012 (2.52%) and *twostage* protocol for iPSYCH2015i (1.83%) and highest when using the *intersection* protocol (iPSYCH2012: 5.89%; iPSYCH2015i: 5.72%). There appears to be a minor gain in variance explained, when using imputed dosages, rather than best guess genotypes for PGS, which is most pronounced when using the *intersection* protocol (iPSYCH2012: 1.54%; iPSYCH2015i: 1.88%). Our results are in line with findings from animal breeding studies^24^ demonstrating reduced genetic prediction accuracy introduced by imputations.

**Figure 4.**
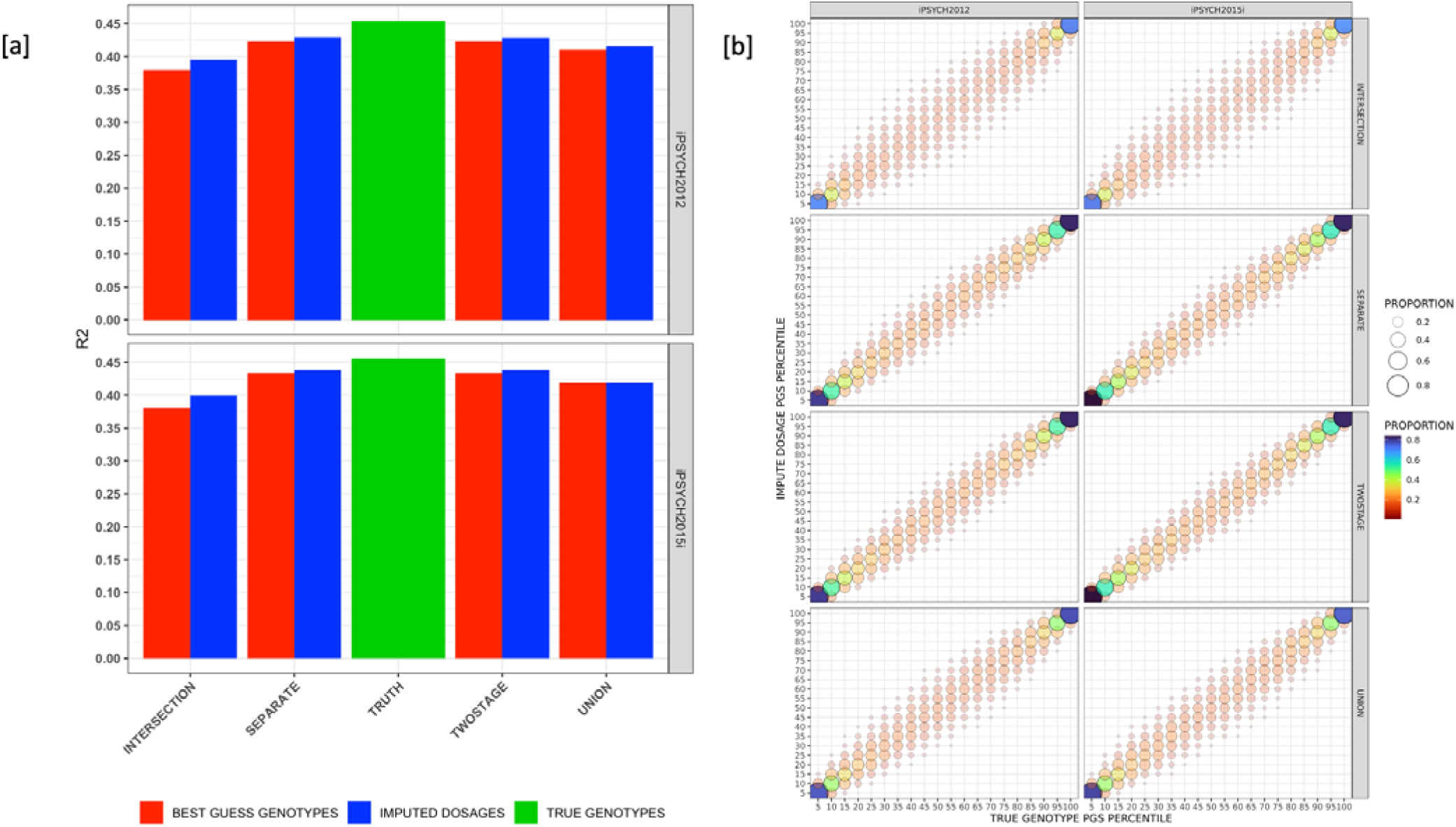
The effects of polygenic scores are attenuated when using imputed data. [a] shows the variance explained (r^2^) in a simulated continuous phenotype with a SNP heritability of 0.5 and the 10,000 SNPs common to both genotyping arrays, masked prior to phasing as causal loci. Variance explained was calculated using the true genotypes, along with imputed dosages and best guess genotypes from the four different data integration protocols. Haplotypes were phased using BEAGLE5 phase-states=560, imputed using BEAGLE5.1 with the HRCv1.1 as the reference. [b] Shows the proportion of individuals in common within each 5-percentile bin when ranked using PGS calculated using true genotypes and imputed dosages.

Another application of PGS is to prioritize individuals in top quantiles of a PGS distribution for monitoring and intervention. To investigate the effect of imputation accuracy on such applications, individuals in iPSYCH2012 and 2015i were grouped into percentiles of PGS risk for the simulated trait based on PGS calculated using genotypes or imputed dosages from the four data integration protocols. The results (Figure 4b) are consistent with prior work^25^ showing a discrepancy in individual rank that is higher in the middle percentiles and lower in the more actionable top percentiles of the PGS distribution. The discordance in individuals in the top percentiles between PGS constructed by true genotypes and imputed dosages is, however, much higher than the 5% previously reported. The overlap in the proportion of individuals ranked in the top 5 percentiles of PGS using true genotypes and imputed dosages is highest in both cohorts when employing either the *separate* or *twostage* protocol (iPSYCH2012: 80%, iPSYCH2015i: 84%) and lowest when using the *intersection* protocol (iPSYCH2012: 71%, iPSYCH2015i: 73%). This overlap in the top 5 percentiles is also greater within iPSYCH2015i, as compared to iPSYCH2012, continuing the trend that higher phasing and imputation accuracies in target samples attributable to higher array density carries through to PGS performance.

### Batch effects

Association studies were performed comparing genotypes and imputed dosages at the masked SNPs from all four data integration protocols in unrelated controls of iPSYCH2012 and 2015i of a homogenous genetic origin with the genotyping array as the outcome (see Methods). The resulting genomic inflation factor in test statistics across different thresholds for imputation quality is shown in figure 5a, supplementary S8. The number of SNPs used in the association tests at each imputation quality threshold is shown in figure 5b.

**Figure 5.**
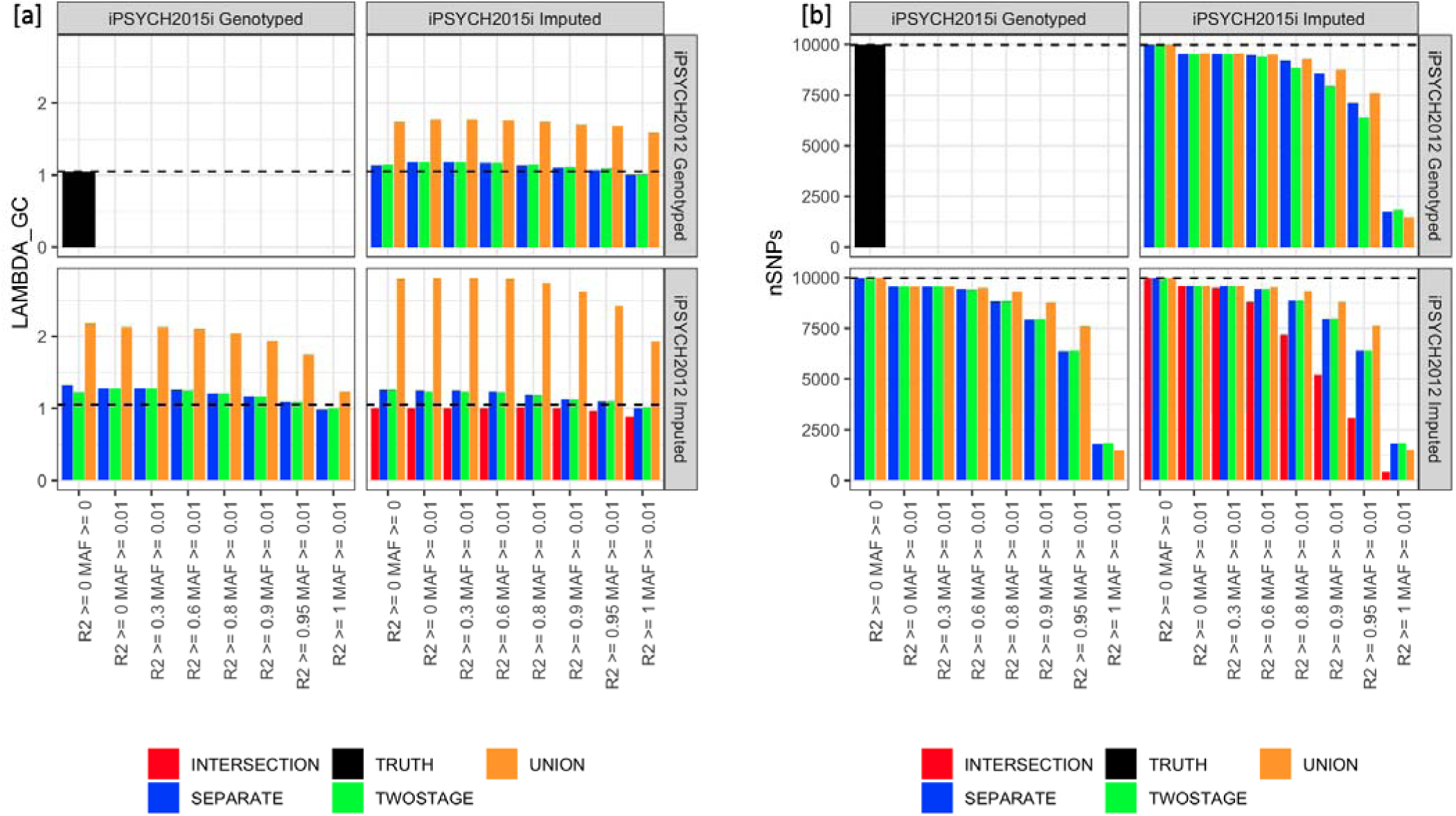
Inflation of test-statistics shows type-I errors associated with imputation. [a] Shows the inflation in test statistics represented using lambda genomic control, when performing an association test at each of the 10,000 SNPs common to both genotyping arrays masked prior to phasing. Controls of a homogeneous genetic origin were compared between the iPSYCH2012 and iPSYCH2015i cohorts with the genotyping array as the outcome at different thresholds of post-imputation quality control across the four different data integration protocols. The dotted horizontal line indicates the baseline ⍰_gc_ when the association test was performed using true genotypes from both arrays. Haplotypes were phased using BEAGLE5 phase-states=560, imputations were done using BEAGLE5.1 with the HRCv1.1 as the reference. [b] Shows the number of SNPs left after each threshold of post imputation quality control across the four data integration protocols.

The baseline for the inflation observed by comparing the genotyped SNPs in controls is 1_gc_ = 1.05. No inflation is observed when comparing SNPs imputed in both iPSYCH2012 and iPSYCH2015i using the *intersection* protocol, while test statistics are most inflated when using the *union* protocol. Using the *separate* and *twostage* protocols, inflation is reduced at high thresholds of BEAGLE imputation r^2^, but not eliminated. For example, in the *Separate* protocol, with SNP imputation quality filter, DR2 >= 0.9, the 1_gc_ = 1.13 when comparing SNPs genotyped in iPSYCH2012 to SNPs imputed in iPSYCH2015i, and 1_gc_ = 1.18 when comparing SNPs imputed in iPSYCH2012 to SNPs genotyped in iPSYCH2015i and 1_gc_ = 1.1 when comparing SNPs imputed in both. At this threshold, 22% of the imputed SNPs are excluded. This analysis suggests that imputations performed from different genotyped backbones, which result in genotyped SNPs being compared to imputed SNPs, will contain batch artifacts that can be difficult to remove by standard SNP exclusion, which might also be complicated, due to a lack of robustness of imputation quality metrics under different data integration protocols (Supplementary S5, Supplementary Figure S3).

## Discussion

As the cost of genotyping drops, the burden of complex trait analysis is moving away from genotyping requisite participants and towards storage, computational requirements, and the bioinformatics expertise to integrate and analyze such datasets^47^. Phasing and imputation have somewhat remained a black box in bioinformatics pipelines with researchers having the opportunity to avail themselves of services like the Michigan imputation server.^48^ to reduce the computational burden of data preparation. However, privacy stipulations governing datasets generated through national biobanks might prohibit use of such services. The benchmarking work presented in this study stresses the importance of making an educated choice of data integration protocols that could introduce a tradeoff among peculiarities such as a sparse marker set, small sample size, high missingness in the input dataset, or the potential of batch artifacts.

The benchmarking of imputation accuracy presented in this study replicates previous findings^23^, suggesting imputation from the intersection of markers when incorporating samples genotyped on multiple arrays leads to a loss of accuracy while imputation from the union of the markers leads to spurious associations with genotyping arrays^23^. Consistent with our hypothesis that the phasing accuracy could be improved by increasing the target sample size by jointly analyzing the two cohorts (by either the union or two-stage protocol) we did observe a drop in SER. However, these improvements did not result in improvements in imputation accuracy, likely reflecting that the phasing tools were not developed with this type of systematic missingness in mind. Until software that can leverage this apparent potential for improvement in phasing accuracy are available, our results suggest that phasing and imputing separately results in equivalent or better imputation accuracy. The higher phasing and imputation accuracies, PGS performance in the sub-cohort of iPSYCH individuals genotyped using the Illumina Global Screening Array, enriched with more common markers as compared to the sub-cohort imputed using the Infinium Psych Array, enriched for rare markers with prior associations to psychiatric phenotypes, suggests that when faced with a choice, it might be more beneficial to prioritize genotyping arrays with more common markers that overlap more with the content of haplotype reference panels. Analysis pipelines and methods focusing on common disease research, rely on established high quality SNP sets, such as HapMap3 and use thresholds to exclude rare markers during QC, effectively rendering them useless for such applications.

Imputed data will contain non-random errors, especially in presence of systemic missingness, as can be the case when genotyping of samples is performed in batches and over time. Therefore, it is critical to consider the sensitivity of any analysis performed on these datasets. Technical artifacts in the genotype generation process are one of the sources of poor performance of PGS across cohorts^27^. While the attenuation introduced in PGS performance and the discordance of individual rank in different percentiles of the risk distribution when PGS are calculated using imputed data as compared to genotyped SNPs has received attention in animal breeding studies, this remains under-researched in human populations. As one of the clinically informative uses of PGS lies in selecting a subset of individuals in the actionable risk percentiles of a PGS distribution^26, 27^, errors introduced during phasing and imputation could have a sizable impact on genetic risk profiling - especially when data is acquired over time and according to different protocols. The presence of spurious associations with genotyping arrays when comparing allele frequencies of genotypes and imputed dosages between cohorts as demonstrated in this study shows the need to pick stringent quality control thresholds for GWAS. As stringent filtering might reduce the power due to exclusion of many imputed SNPs, other approaches such as including the genotyping array as a covariate in regression models or as a fixed effect in linear mixed models need to be further investigated.

Haplotype reference panels employed for phasing and imputation are skewed towards Europeans and the evaluation of imputation accuracy within iPSYCH individuals, grouped by parental birthplace shows differentially worse accuracy in non-Europeans, stressing the need for reference panels with a more genetically diverse catalog of haplotypes, if genotyping arrays and imputation are to be used in precision medicine initiatives in a fair and equitable manner^45, 46^. While considerable attention has been paid to the lack of PGS portability between populations due to less informative SNP effects, less attention has been paid to imputation quality in non-European populations, which introduces an additional source of error, not only in PGS but also in GWAS within these populations. While our comparisons held the reference population constant to the largest set of haplotypes that are currently publicly available, testing the imputation performance with varying references would also be informative. There has been demonstrable improvement in imputation accuracy for individuals of Hispanic/latin and African descent using the NHLBI Trans-Omics for Precision Medicine whole genome sequenced reference panel^49^, but it is currently only available through an imputation server, rendering its usage prohibitive for studies with data privacy stipulations.

In conclusion, this study demonstrates four different ways of integrating data genotyped on multiple arrays with sparse marker overlap. Care should be applied when integrating data sets and building biobanks for precision medicine initiatives, as improper treatment can hurt PGS performance, introduce batch artifacts, and produce systematically lower quality data in non-European samples.

## ACKNOWLEDGEMENTS

Vivek Appadurai is supported by the Lundbeck Foundation post-doctoral grant: R380-2021-1465. Andrew Schork is supported by the Lundbeck Foundation fellowship: R335-2019-2318. The iPSYCH team was supported by grants from the Lundbeck Foundation (R102-A9118, R155-2014-1724, and R248-2017-2003), NIMH (1R01MH124851-01 to A.D.B.) and the Universities and University Hospitals of Aarhus and Copenhagen. The Danish National Biobank resource was supported by the Novo Nordisk Foundation. High-performance computer capacity for handling and statistical analysis of iPSYCH data on the GenomeDK HPC facility was provided by the Center for Genomics and Personalized Medicine and the Centre for Integrative Sequencing, iSEQ, Aarhus University, Denmark (grant to A.D.B.).

## AUTHOR CONTRIBUTIONS

V.A., A.J.S., O.D., A.B. and A.I. conceived the idea and designed the experiments. J.G. performed the genotyping of iPSYCH and QC. V.A., O.D., A.R. performed the data cleaning, bioinformatics, and statistical analyses. V.A., A.J.S. and M.K. wrote the manuscript with participation of all authors. P.B.M., A.D.B., T.W., O.M., M.N., D.H. conceived iPSYCH and obtained funding. All authors read and acknowledged the manuscript.

## SUPPLEMENTARY INFORMATION

## S1. QUALITY CONTROL OF GENOTYPING DATA

The quality control steps prior to phasing are divided into two stages. An initial SNP level QC and a second sample level QC performed on a subset of individuals of a relatively homogenous genetic origin, as determined through the Danish birth registers and principal components analysis, within the iPSYCH sample.

### S1.1. Identifying a genetically homogenous sample subset for QC

Certain steps in the quality control process such as tests of Hardy Weinberg equilibrium, identification of samples with abnormal heterozygosity etc. could be biased by genetic diversity in the dataset. To perform these quality control steps in an unbiased manner; we identify a set of samples of a homogenous genetic origin. To do this, the variant calls from the 1000 genomes phase 3 project^1^ were downloaded in VCF format.

Within each sub-population of the 1000 genomes dataset, we excluded variants for the following reasons:

⍰ Less than 5% minor allele frequency
⍰ Hardy Weinberg p < 10-6
⍰ Pairwise r2 > 0.1 in a 1kB region
⍰ No overlap with the marker set in the Infinium Psych Chip v1.0 and the Global Screening array v2.0.
⍰ Insertions/Deletions
⍰ Regions with extended linkage disequilibrium^2^.

The resulting data was merged with iPSYCH2012 and iPSYCH2015i using PLINK^3^. We performed a principal component analysis using the smartpca module of the eigensoft software package^4^, the principal components were computed using the 1000 genomes samples and the iPSYCH2012, iPSYCH2015i samples were projected into the resulting principal component space.

We further utilized the Danish national birth records to identify a set of 47,586 individuals whose parents and both sets of grandparents were born in Denmark. For each sample in our dataset, we calculate the mahalanobis distance of the sample from the multivariate mean of the joint distribution of the first ten principal components obtained from the 47,586 individuals previously identified. We exclude a sample as an outlier if the distance has a probability less than 5.73x10^-7^ under a chi-square distribution with 10 degrees of freedom. This resulted in 120,890 samples classified as inliers to be used for quality control.

### S1.2. SNP QC

#### S1.2.1 Aligning to the reference

All 26 waves of iPSYCH2012, 78 waves of iPSYCH2015i, Trios2012, Trios2015 and the PGP-UK samples were aligned to Haplotype Reference Consortium v1.1 (hereafter referred to as HRC) using GenotypeHarmonizer v1.4.20-SNAPSHOT^5^. SNP IDs in the target datasets were harmonized to the SNP IDs in the HRC where a match was found, A/T and G/C SNPs were rescued where possible using linkage disequilibrium information, variants absent in the reference, multi-allelic SNPs and indels were excluded.

#### S1.2.2 SNP Missingness

Per SNP and sample missingness were calculated using PLINK 2.0. Genotyping for iPSYCH2012 was performed in 26 waves. We initially excluded variants missing in > 5% of samples in each individual wave. The samples were further merged and variants that were either not genotyped in all 26 waves or were found to be missing in >= 5% of samples in the merged dataset were further excluded. 344,498 SNPs pass this QC.

The genotyping for iPSYCH2015i was performed in 78 waves. We excluded SNPs missing in more than 5% of the samples in each genotyping wave. Samples were merged across batches and SNPs missing in more than 5% of samples across the entire cohort were removed. A total of 558,013 SNPs pass missingness filters.

#### S1.2.3 Differential Missingness between Cases and Controls

We test for SNPs showing differential missingness between cases and controls of a homogenous genetic origin as described in section 1 using the --test-missing option in PLINK. We excluded SNPs that show evidence for differential missingness with an FDR adjusted p-value <= 0.2. 342,837 SNPs in iPSYCH2012 and 555,131 SNPs in iPSYCH2015i pass this filter.

#### S1.2.4 Test of Hardy Weinberg Equilibrium in Controls

The individuals of a homogenous genetic origin as derived in section 1 were further subset to include individuals without any disease diagnosis as ascertained from the Danish national patient registers and a test for Hardy Weinberg equilibrium was performed using the --hardy option in PLINK. We exclude SNPs that fail this test with an FDR adjusted p <= 0.2. 338,104 SNPs in iPSYCH2012 and 544,308 SNPs in iPSYCH2015i pass this QC.

#### S1.2.5 SNPs significantly associated with a genotyping wave or batch

Due to the large sample size of iPSYCH, the genotyping for iPSYCH2012 was performed in 26 waves and the genotyping for iPSYCH2015i was performed in 78 waves. To identify markers showing significant batch effects, we performed 26 and 78 logistic regressions in iPSYCH2012 and iPSYCH2015i respectively where samples of a homogenous genetic origin in a particular wave are cases and samples in other waves are controls. For each SNP, we take the minimum of p-values from all association tests.

The p-values thus selected do not follow a uniform distribution and the cumulative distribution function of drawing minimums from n independent distributions Y = min(p_1_, p_2_, .. p_n_) is given by

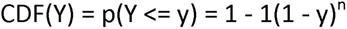

If p_i_ is the i^th^ element in a set of m sorted p-values, the CDF of p_i_ is given by i/m. The i^th^ element in a set of m sorted minimum p-values is given by

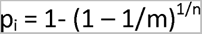

The qq-plot of observed vs expected p-values using the above theoretical distribution suggests some inflation.

FDR adjustment using the above CDF is given by

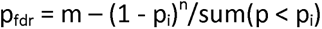

We chose an FDR adjusted p-value cut-off of 0.1 to exclude SNPs, which corresponded to a p-value of 6.31x10^-5^ in iPSYCH2012 and 2.38x10^-6^ in iPSYCH 2016. SNPs passing QC filters, iPSYCH 2012 = 333,308, iPSYCH2015i = 543,422.

#### S1.2.6 Minor Allele Frequency

A subset of 34,545 individuals in iPSYCH2012 were exome sequenced using the Illumina capture kit on HiSeq machines. Quality control was performed using HAIL and variant calling was performed in accordance with the GATK best practices. More details on the data processing have previously been described^6^.

For these individuals, we calculated genotype concordance between the exome sequencing data and genotypes from the iPSYCH2012 array data using bcftools^7^ as shown in supplementary table 1. We observe that the concordance between genotyped and next generation sequencing datasets drops sharply at minor allele frequencies below 0.001. So, we chose this as a sensible threshold for censoring SNPs. SNPs passing QC filters: iPSYCH2012: 261,551, iPSYCH2015i: 460,445.

**Supplementary Table 1.** Concordance between genotypes from Infinium Psych Chip v1.0 and whole exome sequencing data in a subset of 34,545 individuals in iPSYCH2012.

| Allele Frequency Bin | Concordance between genotyping array and Exome Sequencing Data | Number of SNPs |
| --- | --- | --- |
| 0.00001 - 0.0001 | 0.4085 | 20701 |
| 0.0001 - 0.001 | 0.7976 | 30367 |
| 0.001 - 0.01 | 0.9676 | 14145 |
| 0.01 - 0.1 | 0.9966 | 6795 |
| 0.1 - 0.5 | 0.999 | 5081 |
| 0.5 - 1 | 0.9991 | 28 |

#### S1.2.7 SNP Masking

To evaluate the performance of missing data imputation, we randomly selected 10,000 SNPs that were genotyped on both the Illumina PsychArray v1.0 and the Illumina Global Screening Array v2.0 using the sample function in R. These were excluded prior to haplotype estimation. SNPs used for haplotype estimation and imputation, iPSYCH2012: 251,551, iPSYCH2015i: 450,445.

### S1.3. SAMPLE QC

#### S1.3.1 Abnormal Heterozygosity

Abnormal levels of heterozygosity that cannot adequately be explained by admixture, population structure or runs of homozygosity could indicate sample contamination. To identify individuals with heterozygosity that cannot be accounted for by population phenomena, we use an approach described by the UK biobank (https://biobank.ctsu.ox.ac.uk/crystal/crystal/docs/genotyping_qc.pdf). Per sample heterozygosity, homozygosity and missingness were calculated using PLINK --het, --homozyg and -- missing options respectively. Ancestry adjusted heterozygosity is computed as the residuals from the model shown below:

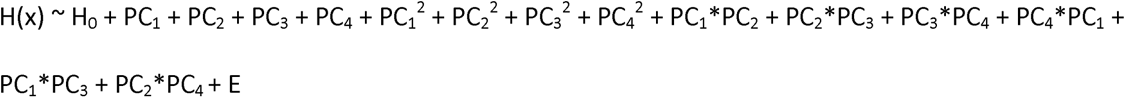

Where H(x) = Observed heterozygosity

H_0_ = Mean heterozygosity/Intercept

PC_1_, PC_2_, PC_3_, PC_4_ = First four principal components of genetic ancestry

E = Residual/Ancestry adjusted heterozygosity

We further fit two linear models predicting the observed and ancestry adjusted heterozygosities from runs of homozygosity calculated using PLINK. Samples are flagged as outliers if the observed and ancestry adjusted heterozygosity as well as the residuals from the models fit against runs of homozygosity are four standard deviations away from the mean. 166 samples from iPSYCH2012 and 98 samples from iPSYCH2015i failed this quality check and were excluded.

#### S1.3.2 Sample Duplication

A total of 121 samples were found to be genotyped more than once across the 26 waves in iPSYCH2012. Further, mapping sample identifiers to unique identifiers from the registers yielded 159 sample identifiers in iPSYCH2012 and 25 sample identifiers in iPSYCH2015i mapping to a non-unique identifier in the registry. Two samples from iPSYCH2012 were found to be genotyped again in iPSYCH2015i due to the randomness of ascertainment. In each case, the sample with lower missingness was retained. 6 samples in iPSYCH2012 and 1 sample in iPSYCH2015i were genotyped as part of the trios and were excluded.

Kinship analysis performed using KING^8^ revealed three monozygotic twins in iPSYCH2012 and ten monozygotic twins in iPSYCH2015i. In each case, the case was retained and if both samples were cases, the sample with higher missingness was excluded.

#### S1.3.3 Sample Missingness

Two samples from the iPSYCH2012 cohort were excluded for excessive missingness (> 5%).

This left us with 80,876 samples in iPSYCH2012, genotyped at 251,551 loci and 48,974 individuals in iPSYCH2015i, genotyped at 450,445 loci to be used as a backbone for haplotype estimation and missing data imputation.

## S2. ANCESTRY COMPOSITION OF iPSYCH

**Supplementary Table 2.** Ancestry composition of iPSYCH by parental birthplace as obtained from the Danish Civil Registers^9^. Underscore delimited combinations indicate parents born in different regions.

| Parental Birthplace | iPSYCH2012 | iPSYCH2015i |
| --- | --- | --- |
| Denmark | 67044 | 41673 |
| Denmark_Europe | 2416 | 1591 |
| Denmark_Scandinavia | 1476 | 913 |
| Europe | 1169 | 785 |
| Denmark_Unknown | 829 | 543 |
| MiddleEast | 775 | 563 |
| Asia | 594 | 384 |
| Asia_Denmark | 581 | 292 |
| Africa | 473 | 277 |
| Denmark_Greenland | 435 | 284 |
| Africa_Denmark | 431 | 292 |
| Denmark_NorthAmerica | 363 | 234 |
| Denmark_MiddleEast | 354 | 216 |
| Denmark_SouthAmerica | 235 | 184 |
| Scandinavia | 109 | 67 |

## S3. SWITCH ERROR RATES

**Supplementary Table 3.**
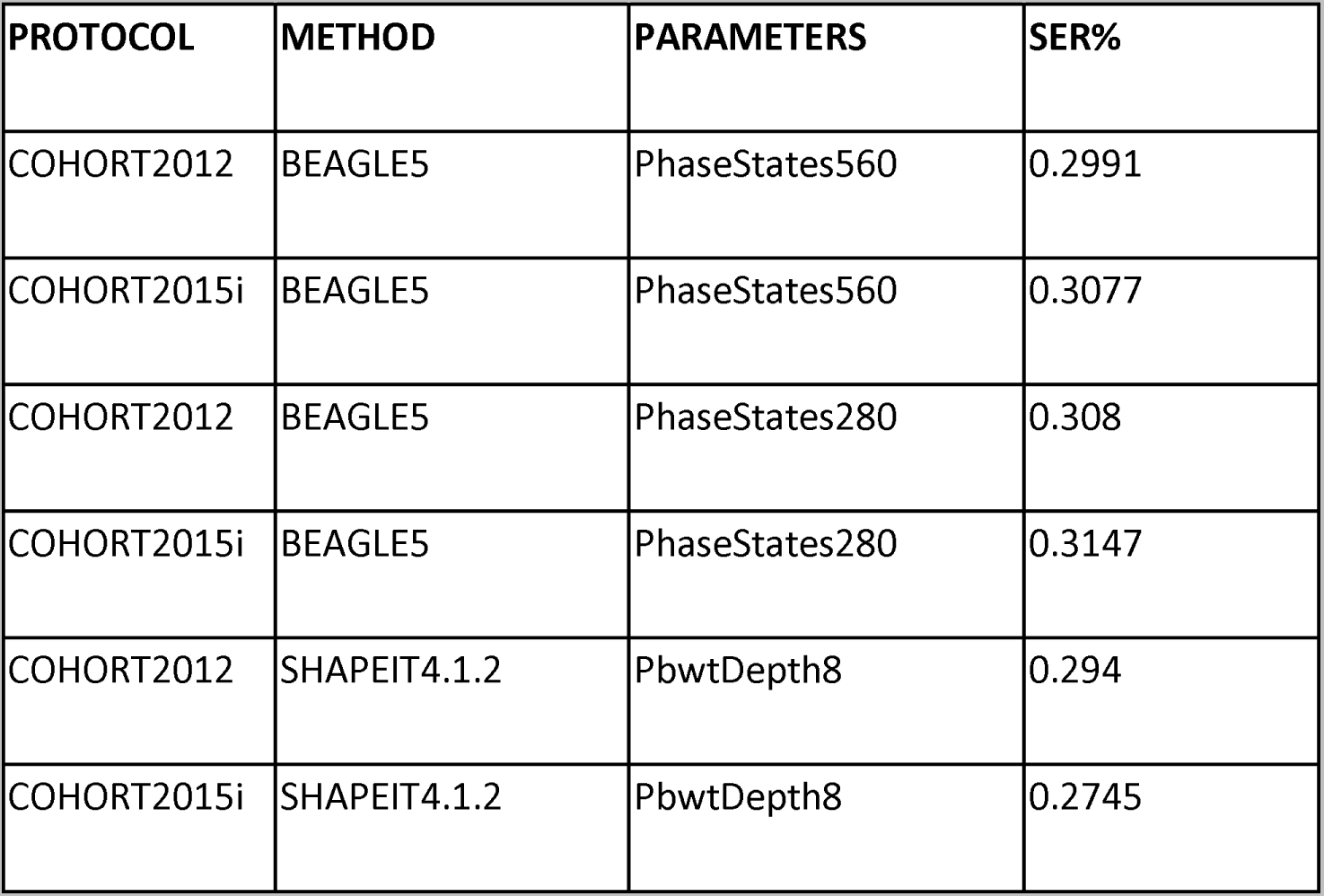
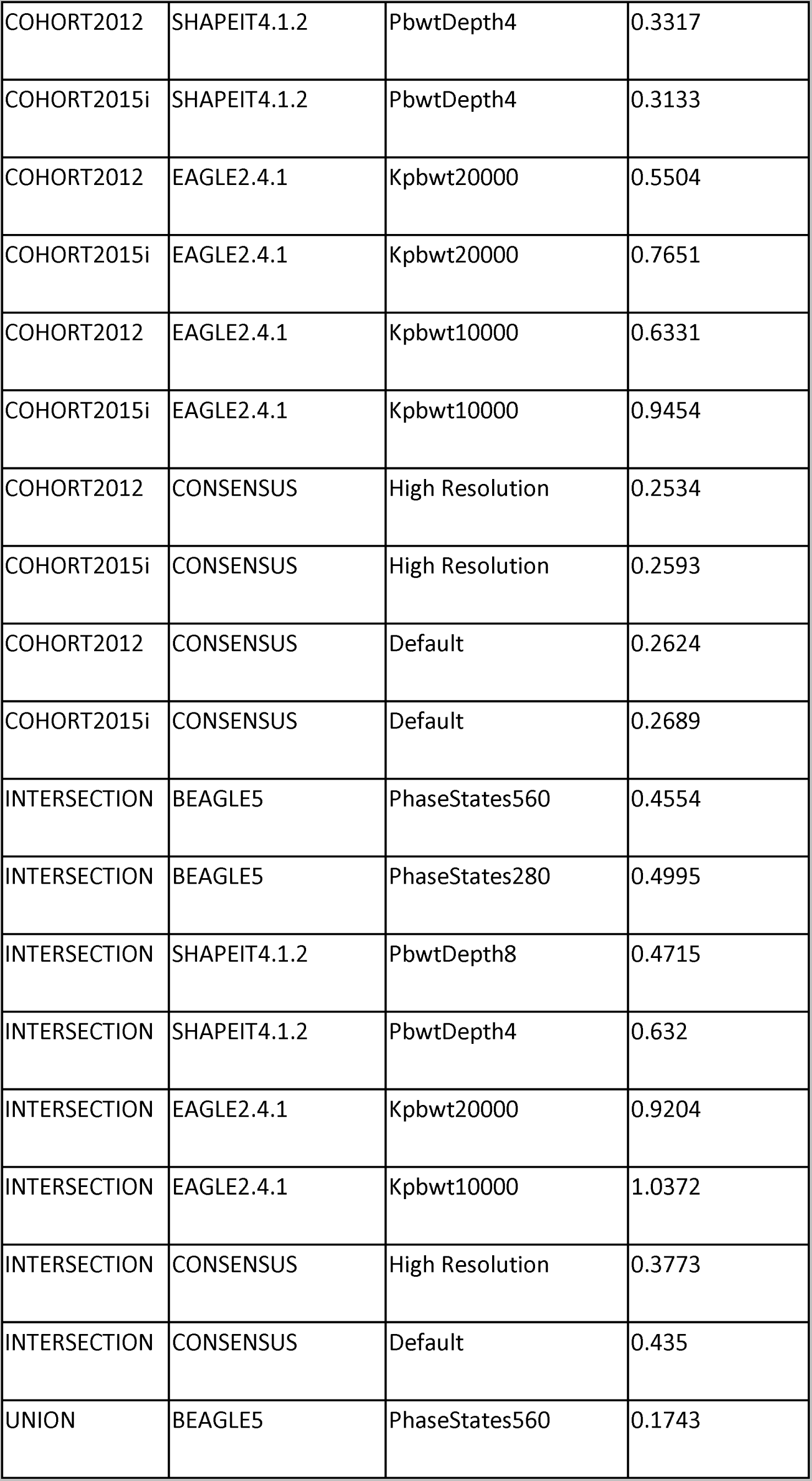
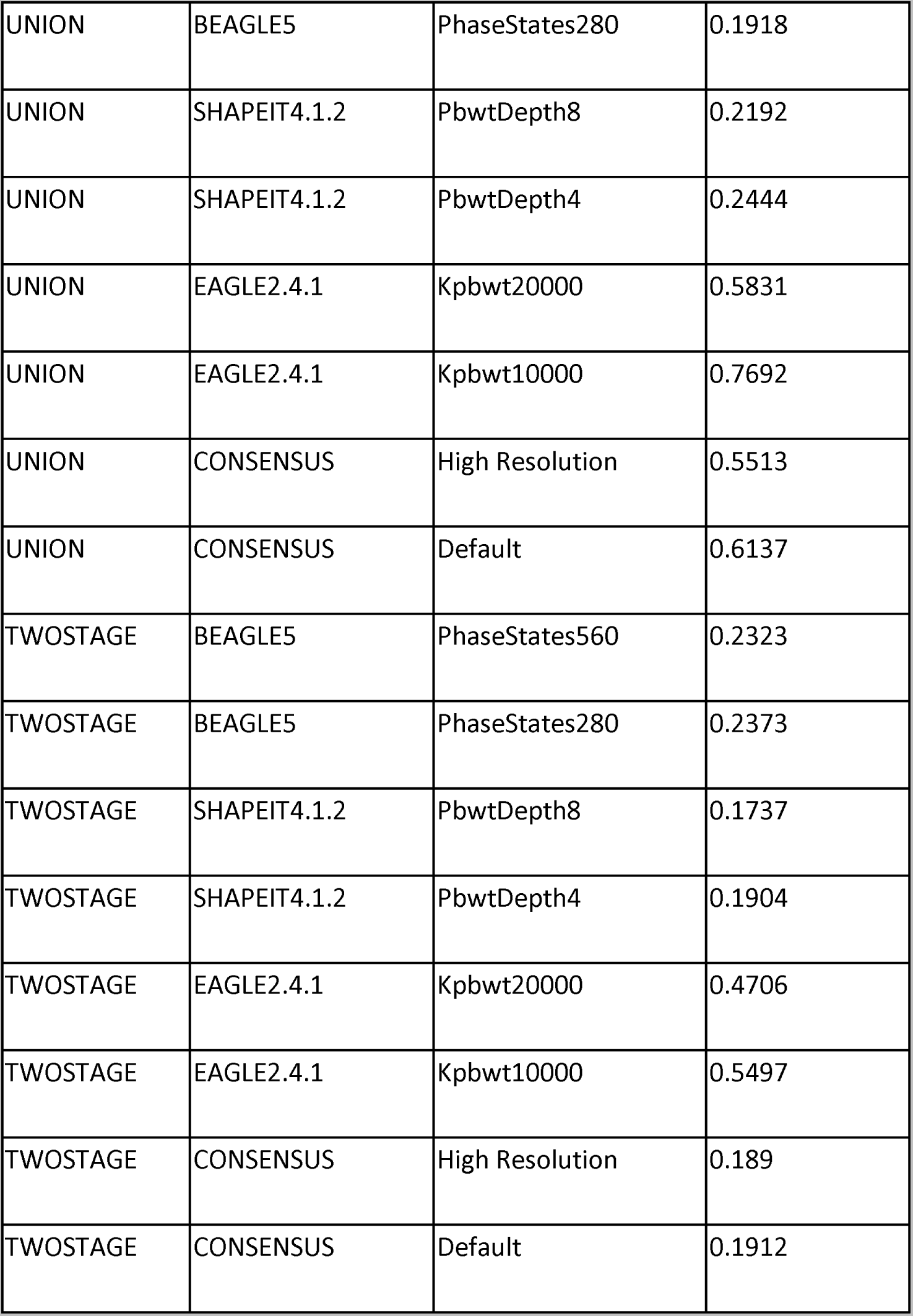
Phasing accuracy as indicated by switch error rates obtained by comparing the mendelian transmission of phase to computationally estimated phase within 124 trio offspring for whom parental genotypes are known at heterozygous loci genotyped on both iPSYCH arrays.

The marker coverage from the iPSYCH genotyping arrays is not uniform across all chromosomes. As shown in supplementary figure 1, this leads to a variability in the accuracy of haplotype estimation by chromosome number.

**Supplementary figure 1.**
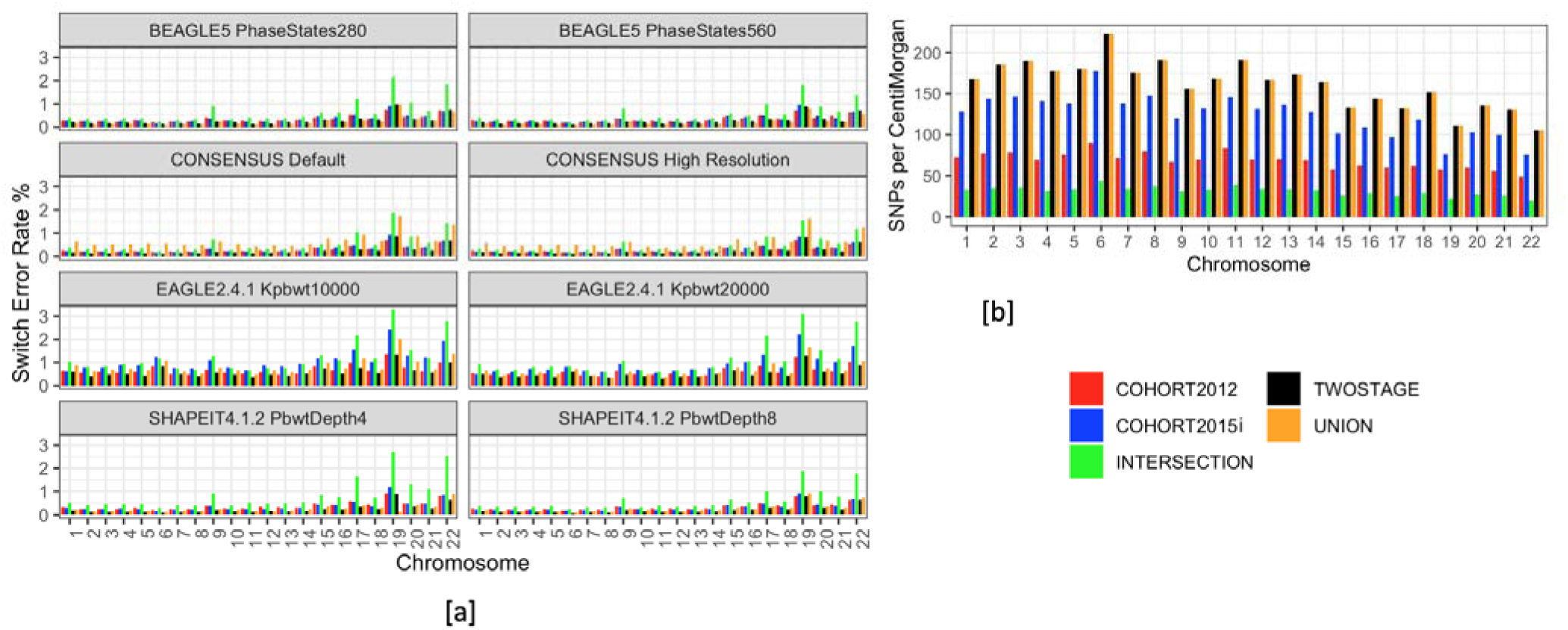
[a] Haplotype estimation accuracy as shown by switch error rates obtained from comparing computationally assigned phase to mendelian transmission in 124 trio offspring whose parental genotypes are known. [b] SNP density across chromosomes within each data integration protocol.

## S4. IMPUTATION ACCURACY WITHIN PERSONAL GENOMES PROJECT - UK SAMPLES

BAM files corresponding to 10 samples from the personal genomes project - UK^10^ were downloaded from the European Genome-Phenome Archive (EGA, study accession: PRJEB17529), sample accessions (SAMEA4545245, SAMEA4545246, SAMEA4545247, SAMEA4545248, SAMEA4545249, SAMEA4545250, SAMEA4545251, SAMEA4545252, SAMEA4545253, SAMEA4545254). Variant calling was performed using samtools mpileup and the samples were further downsampled to each of the two iPSYCH genotyping arrays and added to cohorts arising from each data integration protocol prior to phasing and imputation. The accuracy of the imputation was calculated as the squared Pearson correlation coefficient between the imputed dosages and variant calls at 6.5 million loci not genotyped on either iPSYCH array. The results as shown in supplementary figures 2a, b across minor allele frequency bins as ascertained from the HRCv1.1 haplotype reference panel show similar results to the results obtained by gauging the accuracy at the 10,000 SNPs masked prior to phasing. The accuracy of imputation appears to rely more on choice of data integration protocol than haplotype estimation tool. The haplotypes obtained from SHAPEIT4.1.2 in presence of high missingness introduced by the *union* protocol led to inaccurate imputations.

A comparison of imputation accuracy between the two iPSYCH genotyping arrays as shown in supplementary figure 2c reveals that all tools yield more accurate imputations in the cohort generated using the denser Illumina global screening array v2.0, despite a relatively lesser sample size for haplotype estimation as compared to the cohort generated using the Infinium PsychChip v1.0 with less dense SNP information but a higher sample size.

**Supplementary Figure 2.**
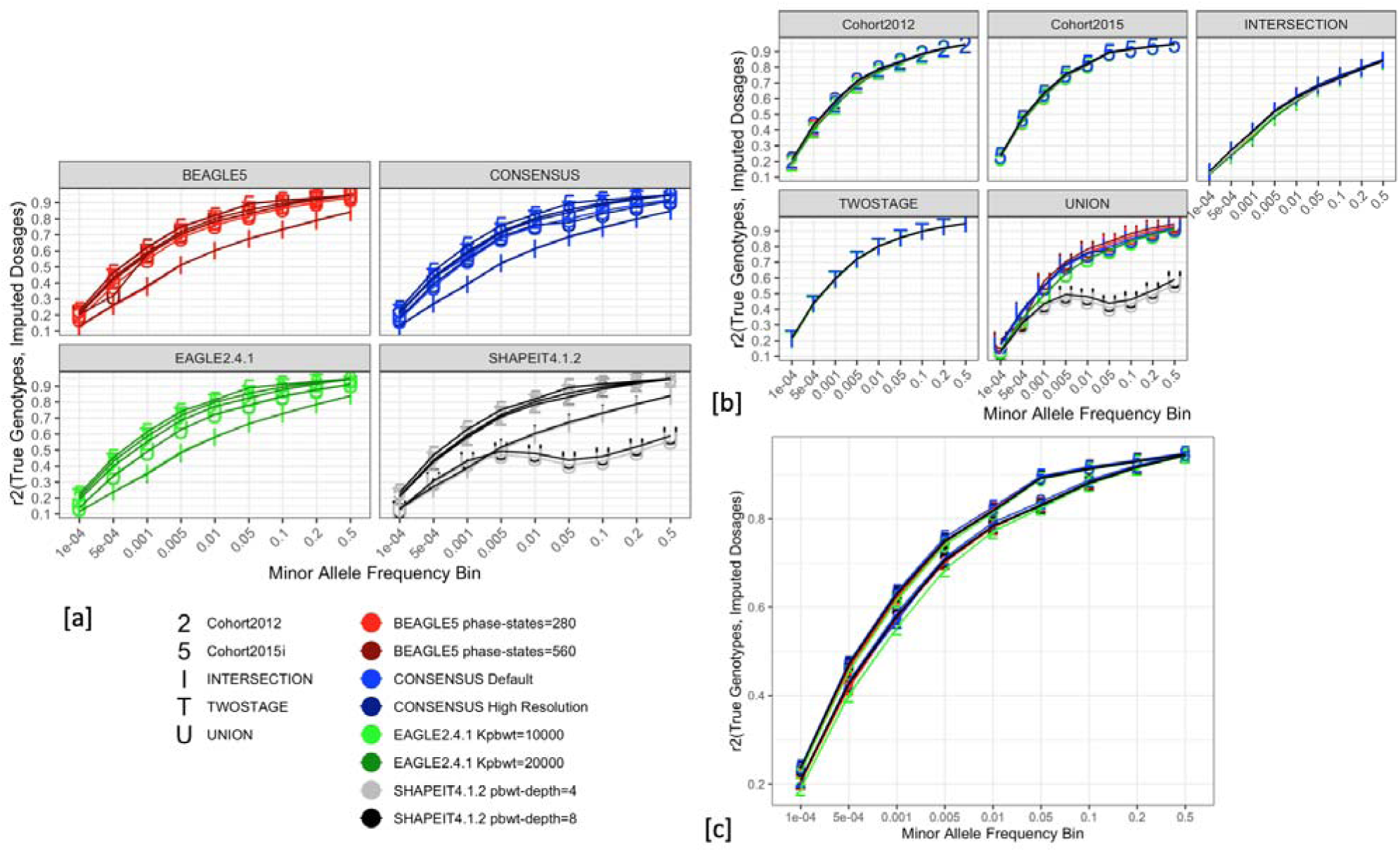
Accuracy of imputation within the personal genomes project - UK whole genome sequenced samples, calculated as the squared Pearson correlation coefficient between imputed dosages and true genotypes at loci absent from either iPSYCH genotyping array. [a] Grouped by choice of haplotype estimation tool. [b] Grouped by choice of data integration protocol. [c] Comparison between imputation accuracy obtained by using each iPSYCH genotyping array.

## S5. RELIABILITY OF IMPUTATION QUALITY METRICS

Imputation software, such as BEAGLE5.1 provides an estimated quality score for imputed dosages (BEAGLE-r^2^) at each SNP, which is a predicted correlation between the true and estimated genotypes at a given variant. The r^2^ at an imputed locus is an important quantity, as it can be used to estimate the reduction in effective sample size for an association test^11^ and as a filtering threshold to ensure only high quality markers are used for association tests and polygenic scoring^12^. We sought to evaluate the robustness of this metric across data integration protocols by comparing it to the empirical imputation accuracy (Empirical-r^2^) calculated from the 10,000 masked SNPs (Supplementary Figure 3). The squared Pearson correlation coefficient of BEAGLE-r^2^ and EMPIRICAL-r^2^is highest for *intersection* protocol (r^2^_BEAGLE-r2, EMPIRICAL-r2_ = 0.98) protocol and lowest for the *union* (r^2^_BEAGLE-r2, EMPIRICAL-r2_ = 0.77) (Supplementary Figure 3). Hence, uncertainties introduced by high genotype missingness in the target dataset, prior to phasing travels through the whole genome imputation pipeline, leading to a potential inclusion of genotype dosages, estimated at less than the recommended thresholds and impacting the accuracy of estimates and replicability of complex trait analyses.

**Supplementary Figure 3.**
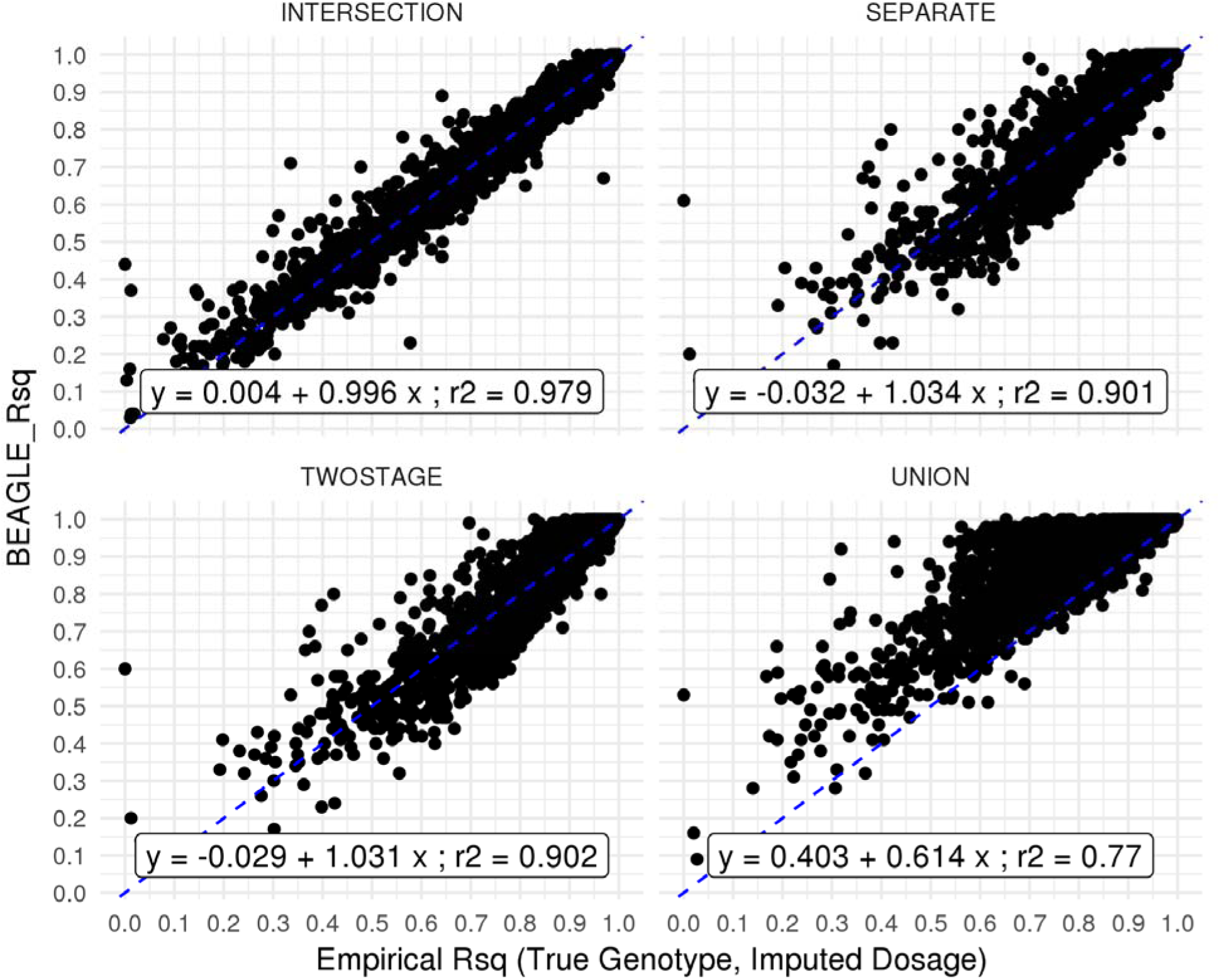
The relationship between empirical imputation accuracy, as measured by the squared Pearson correlation coefficient of true genotypes and imputed dosages at 10,000 masked SNPs, and BEAGLE r^2^ within each data integration protocol. The plot shows the BEAGLE r^2^ is best calibrated for the imputations from the *intersection* protocol whereas it overestimates the accuracy, in presence of high genotype missingness, as present in the *union* protocol.

## S6. IMPUTATION ACCURACY IN NON-EUROPEAN AND ADMIXED SAMPLES ACROSS DATA INTEGRATION PROTOCOLS

**Supplementary Figure 4.**
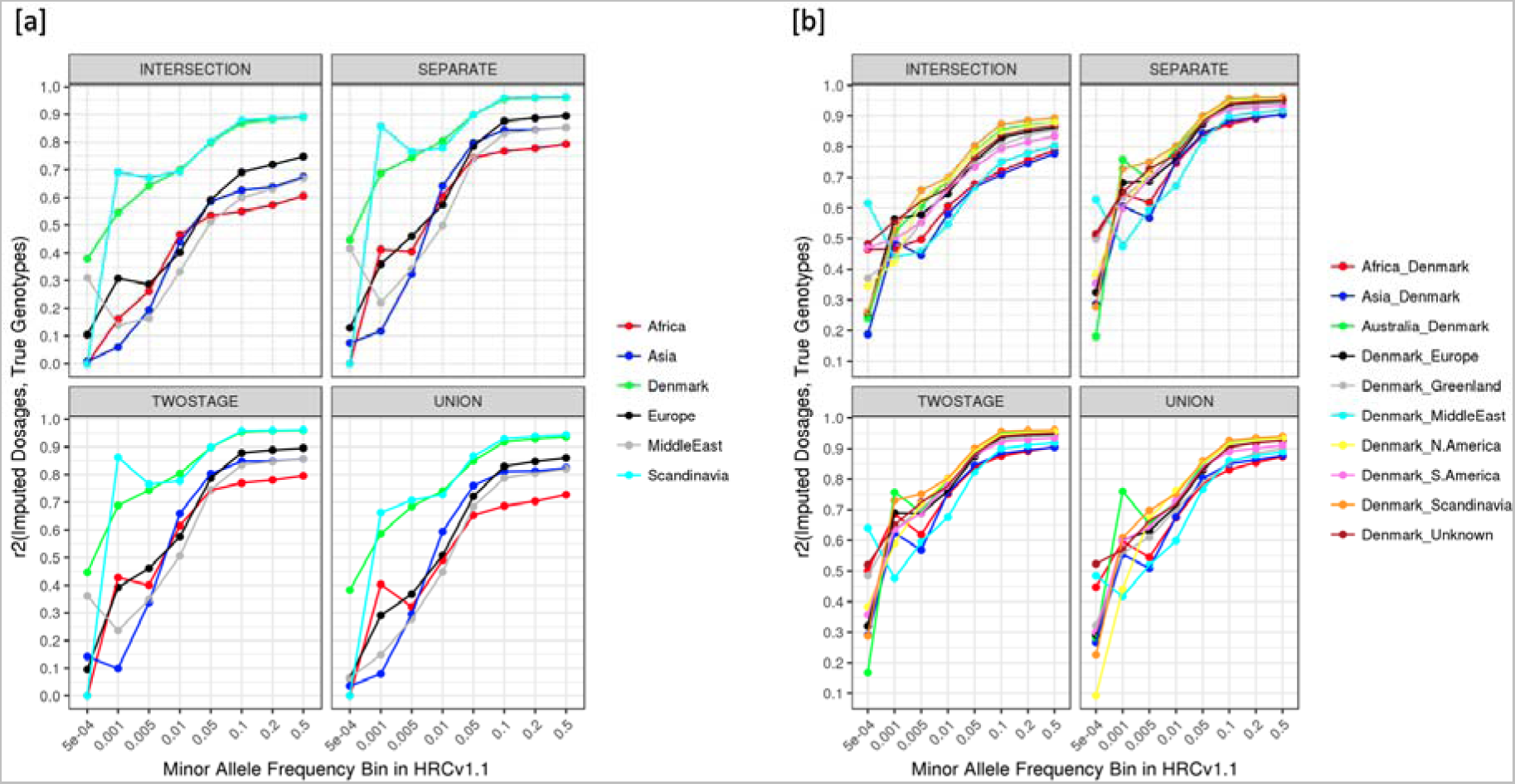
Accuracy of imputation varies by parental origin. The attenuation in imputation accuracy within samples of non-European origin is further magnified by choice of data integration protocol. [a] Shows the accuracy of imputation within the 10,000 masked SNPs at different minor allele frequency bins within samples grouped by the birthplace of their parents according to the Danish civil registers across all four data integration protocols. [b] Shows the accuracy of imputation within the 10,000 masked SNPs within samples where at least one parent was born in Denmark.

## S7. PGS ANALYSIS

**Supplementary Table 4.**
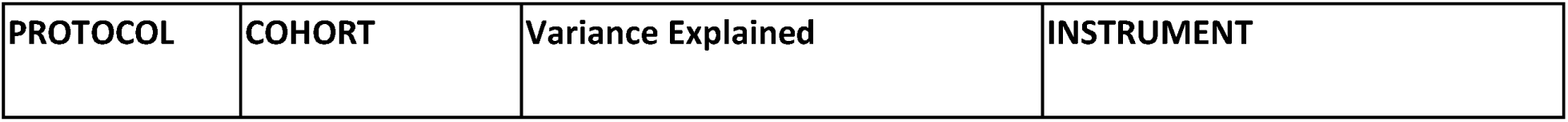
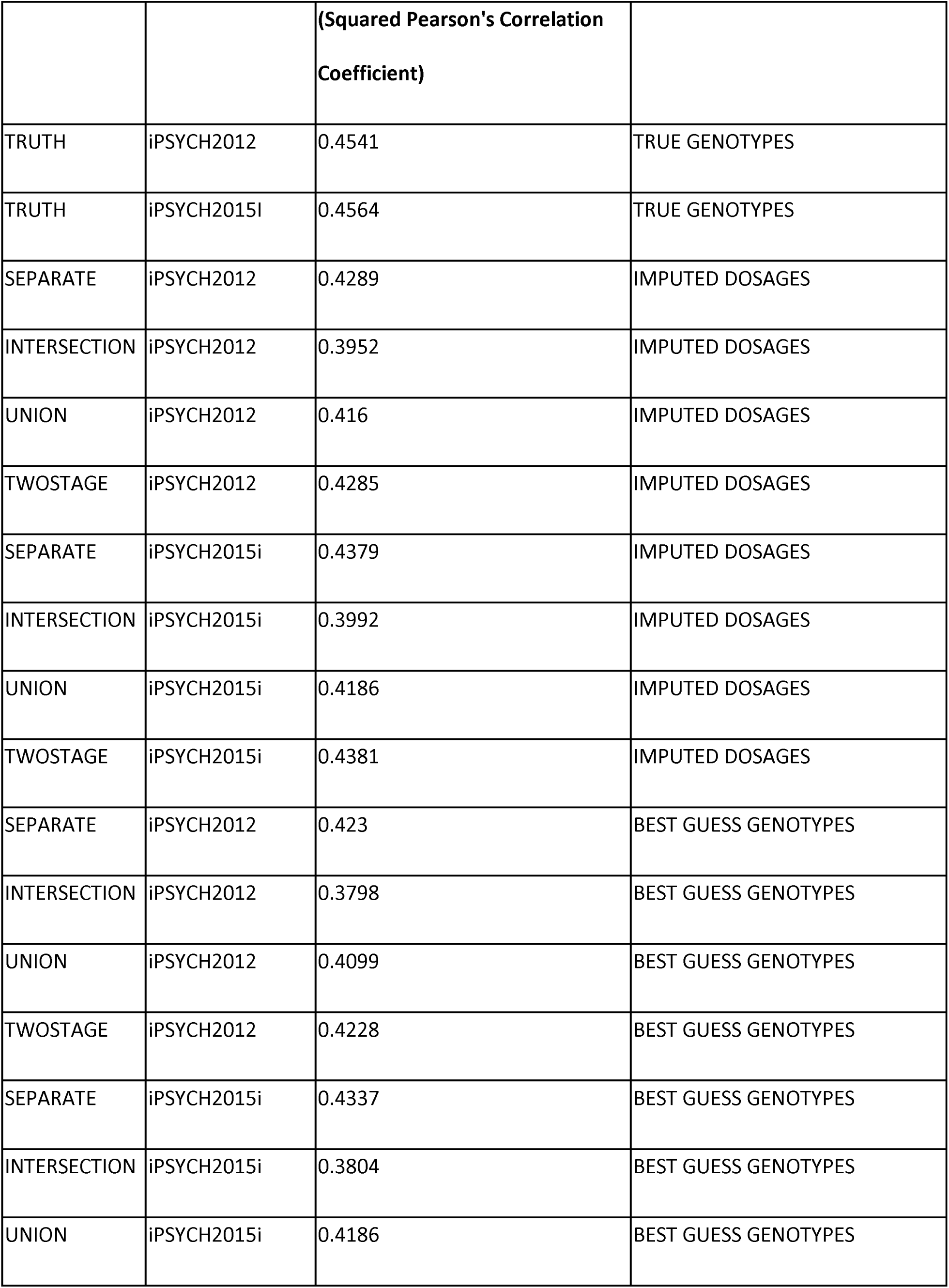
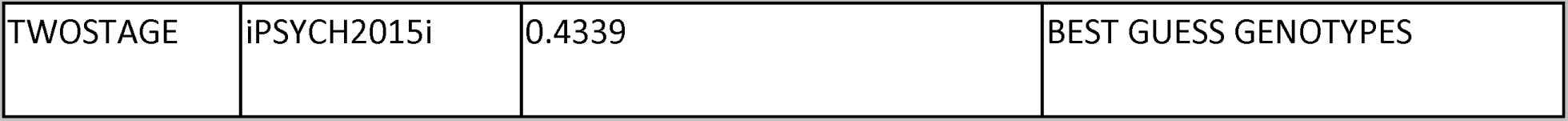
Variance explained in a simulated continuous phenotype with a SNP heritability of 0.5 and the 10,000 masked SNPs as causal loci by true genotypes and imputed dosages, best guess genotypes across the four data integration protocols.

## S8. BATCH EFFECTS

**Supplementary Table 5.** Batch effects, as demonstrated by the inflation in test statistics when performing GWAS with genotyped and imputed dosages from the two iPSYCH cohorts at the 10,000 masked SNPs in controls of European origin with the genotyping array as the outcome.

| PROTOCOL | iPSYCH2012 | iPSYCH2015i | R2 | MAF | N | LAMBDA_GC |
| --- | --- | --- | --- | --- | --- | --- |
| TWOSTAGE | Imputed | Imputed | 0 | 0.01 | 9566 | 1.23 |
| TWOSTAGE | Imputed | Imputed | 0.3 | 0.01 | 9564 | 1.23 |
| TWOSTAGE | Imputed | Imputed | 0.6 | 0.01 | 9430 | 1.22 |
| TWOSTAGE | Imputed | Imputed | 0.8 | 0.01 | 8871 | 1.18 |
| TWOSTAGE | Imputed | Imputed | 0.9 | 0.01 | 7964 | 1.13 |
| TWOSTAGE | Imputed | Imputed | 0.95 | 0.01 | 6410 | 1.1 |
| TWOSTAGE | Imputed | Imputed | 1 | 0.01 | 1845 | 1.01 |
| TWOSTAGE | Genotyped | Imputed | 0 | 0.01 | 9543 | 1.18 |
| TWOSTAGE | Genotyped | Imputed | 0.3 | 0.01 | 9541 | 1.18 |
| TWOSTAGE | Genotyped | Imputed | 0.6 | 0.01 | 9407 | 1.17 |
| TWOSTAGE | Genotyped | Imputed | 0.8 | 0.01 | 8849 | 1.15 |
| TWOSTAGE | Genotyped | Imputed | 0.9 | 0.01 | 7945 | 1.11 |
| TWOSTAGE | Genotyped | Imputed | 0.95 | 0.01 | 6397 | 1.09 |
| TWOSTAGE | Genotyped | Imputed | 1 | 0.01 | 1841 | 1.01 |
| TWOSTAGE | Imputed | Genotyped | 0 | 0.01 | 9543 | 1.28 |
| TWOSTAGE | Imputed | Genotyped | 0.3 | 0.01 | 9541 | 1.28 |
| TWOSTAGE | Imputed | Genotyped | 0.6 | 0.01 | 9407 | 1.25 |
| TWOSTAGE | Imputed | Genotyped | 0.8 | 0.01 | 8849 | 1.21 |
| TWOSTAGE | Imputed | Genotyped | 0.9 | 0.01 | 7945 | 1.16 |
| TWOSTAGE | Imputed | Genotyped | 0.95 | 0.01 | 6397 | 1.09 |
| TWOSTAGE | Imputed | Genotyped | 1 | 0.01 | 1841 | 1 |
| SEPARATE | Imputed | Imputed | 0 | 0.01 | 9569 | 1.25 |
| SEPARATE | Imputed | Imputed | 0.3 | 0.01 | 9566 | 1.25 |
| SEPARATE | Imputed | Imputed | 0.6 | 0.01 | 9431 | 1.23 |
| SEPARATE | Imputed | Imputed | 0.8 | 0.01 | 8869 | 1.19 |
| SEPARATE | Imputed | Imputed | 0.9 | 0.01 | 7957 | 1.13 |
| SEPARATE | Imputed | Imputed | 0.95 | 0.01 | 6404 | 1.1 |
| SEPARATE | Imputed | Imputed | 1 | 0.01 | 1822 | 1 |
| SEPARATE | Genotyped | Imputed | 0 | 0.01 | 9546 | 1.18 |
| SEPARATE | Genotyped | Imputed | 0.3 | 0.01 | 9544 | 1.18 |
| SEPARATE | Genotyped | Imputed | 0.6 | 0.01 | 9497 | 1.17 |
| SEPARATE | Genotyped | Imputed | 0.8 | 0.01 | 9220 | 1.14 |
| SEPARATE | Genotyped | Imputed | 0.9 | 0.01 | 8565 | 1.1 |
| SEPARATE | Genotyped | Imputed | 0.95 | 0.01 | 7105 | 1.07 |
| SEPARATE | Genotyped | Imputed | 1 | 0.01 | 1747 | 1 |
| SEPARATE | Imputed | Genotyped | 0 | 0.01 | 9561 | 1.28 |
| SEPARATE | Imputed | Genotyped | 0.3 | 0.01 | 9559 | 1.28 |
| SEPARATE | Imputed | Genotyped | 0.6 | 0.01 | 9423 | 1.26 |
| SEPARATE | Imputed | Genotyped | 0.8 | 0.01 | 8858 | 1.21 |
| SEPARATE | Imputed | Genotyped | 0.9 | 0.01 | 7939 | 1.16 |
| SEPARATE | Imputed | Genotyped | 0.95 | 0.01 | 6384 | 1.09 |
| SEPARATE | Imputed | Genotyped | 1 | 0.01 | 1792 | 0.98 |
| UNION | Imputed | Imputed | 0 | 0.01 | 9571 | 2.81 |
| UNION | Imputed | Imputed | 0.3 | 0.01 | 9571 | 2.81 |
| UNION | Imputed | Imputed | 0.6 | 0.01 | 9536 | 2.8 |
| UNION | Imputed | Imputed | 0.8 | 0.01 | 9326 | 2.74 |
| UNION | Imputed | Imputed | 0.9 | 0.01 | 8799 | 2.62 |
| UNION | Imputed | Imputed | 0.95 | 0.01 | 7628 | 2.43 |
| UNION | Imputed | Imputed | 1 | 0.01 | 1478 | 1.93 |
| UNION | Genotyped | Imputed | 0 | 0.01 | 9548 | 1.77 |
| UNION | Genotyped | Imputed | 0.3 | 0.01 | 9548 | 1.77 |
| UNION | Genotyped | Imputed | 0.6 | 0.01 | 9513 | 1.76 |
| UNION | Genotyped | Imputed | 0.8 | 0.01 | 9304 | 1.74 |
| UNION | Genotyped | Imputed | 0.9 | 0.01 | 8779 | 1.7 |
| UNION | Genotyped | Imputed | 0.95 | 0.01 | 7612 | 1.68 |
| UNION | Genotyped | Imputed | 1 | 0.01 | 1476 | 1.59 |
| UNION | Imputed | Genotyped | 0 | 0.01 | 9548 | 2.13 |
| UNION | Imputed | Genotyped | 0.3 | 0.01 | 9548 | 2.13 |
| UNION | Imputed | Genotyped | 0.6 | 0.01 | 9513 | 2.11 |
| UNION | Imputed | Genotyped | 0.8 | 0.01 | 9304 | 2.04 |
| UNION | Imputed | Genotyped | 0.9 | 0.01 | 8779 | 1.94 |
| UNION | Imputed | Genotyped | 0.95 | 0.01 | 7612 | 1.75 |
| UNION | Imputed | Genotyped | 1 | 0.01 | 1476 | 1.23 |
| INTERSECTION | Imputed | Imputed | 0 | 0.01 | 9570 | 1 |
| INTERSECTION | Imputed | Imputed | 0.3 | 0.01 | 9499 | 1 |
| INTERSECTION | Imputed | Imputed | 0.6 | 0.01 | 8802 | 1 |
| INTERSECTION | Imputed | Imputed | 0.8 | 0.01 | 7183 | 1.01 |
| INTERSECTION | Imputed | Imputed | 0.9 | 0.01 | 5226 | 1 |
| INTERSECTION | Imputed | Imputed | 0.95 | 0.01 | 3096 | 0.97 |
| INTERSECTION | Imputed | Imputed | 1 | 0.01 | 421 | 0.89 |

* Supplementary tables 6 - 9 in appendix.

## References

1. Visscher, P. M., Brown, M. A., McCarthy, M. I. & Yang, J. Five years of GWAS discovery. Am. J. Hum. Genet. 90, 7–24 (2012).

2. Visscher, P. M. et al. 10 Years of GWAS Discovery: Biology, Function, and Translation. Am. J. Hum. Genet. 101, 5–22 (2017).

3. Marchini, J. & Howie, B. Genotype imputation for genome-wide association studies. Nat. Rev. Genet. 11, 499–511 (2010).

4. Das, S., Abecasis, G. R. & Browning, B. L. Genotype Imputation from Large Reference Panels. Annu. Rev. Genomics Hum. Genet. 19, 73–96 (2018).

5. Li, Y., Willer, C., Sanna, S. & Abecasis, G. Genotype imputation. Annu. Rev. Genomics Hum. Genet. 10, 387–406 (2009).

6. Zeggini, E. & Ioannidis, J. P. A. Meta-analysis in genome-wide association studies. Pharmacogenomics 10, 191–201 (2009).

7. Choi, S. W. & O’Reilly, P. F. PRSice-2: Polygenic Risk Score software for biobank-scale data. Gigascience 8, (2019).

8. Browning, B. L., Zhou, Y. & Browning, S. R. A One-Penny Imputed Genome from Next-Generation Reference Panels. Am. J. Hum. Genet. 103, 338–348 (2018).

9. Delaneau, O., Zagury, J.-F., Robinson, M. R., Marchini, J. L. & Dermitzakis, E. T. Accurate, scalable and integrative haplotype estimation. Nat. Commun. 10, 5436 (2019).

10. Loh, P.-R. et al. Reference-based phasing using the Haplotype Reference Consortium panel. Nat. Genet. 48, 1443–1448 (2016).

11. Li, N. & Stephens, M. Modeling linkage disequilibrium and identifying recombination hotspots using single-nucleotide polymorphism data. Genetics 165, 2213–2233 (2003).

12. Bycroft, C. et al. The UK Biobank resource with deep phenotyping and genomic data. Nature 562, 203–209 (2018).

13. 1000 Genomes Project Consortium et al. A global reference for human genetic variation. Nature 526, 68–74 (2015).

14. Zook, J. M. et al. Extensive sequencing of seven human genomes to characterize benchmark reference materials. Scientific data vol. 3 160025 (2016).

15. Banda, Y. et al. Characterizing Race/Ethnicity and Genetic Ancestry for 100,000 Subjects in the Genetic Epidemiology Research on Adult Health and Aging (GERA) Cohort. Genetics 200, 1285–1295 (2015).

16. Panagiotou, O. A., Willer, C. J., Hirschhorn, J. N. & Ioannidis, J. P. A. The power of meta-analysis in genome-wide association studies. Annu. Rev. Genomics Hum. Genet. 14, 441–465 (2013).

17. Zaitlen, N. & Eskin, E. Imputation aware meta-analysis of genome-wide association studies. Genet. Epidemiol. 34, 537–542 (2010).

18. Browning, S. R. & Browning, B. L. Haplotype phasing: existing methods and new developments. Nat. Rev. Genet. 12, 703–714 (2011).

19. Pedersen, C. B. et al. The iPSYCH 2012 case-cohort sample: new directions for unravelling genetic and environmental architectures of severe mental disorders. Mol. Psychiatry 23, 6–14 (2018).

20. Loh, P.-R., Palamara, P. F. & Price, A. L. Fast and accurate long-range phasing in a UK Biobank cohort. Nat. Genet. 48, 811–816 (2016).

21. Sinnott, J. A. & Kraft, P. Artifact due to differential error when cases and controls are imputed from different platforms. Hum. Genet. 131, 111–119 (2012).

22. Uh, H.-W. et al. How to deal with the early GWAS data when imputing and combining different arrays is necessary. Eur. J. Hum. Genet. 20, 572–576 (2012).

23. Johnson, E. O. et al. Imputation across genotyping arrays for genome-wide association studies: assessment of bias and a correction strategy. Hum. Genet. 132, 509–522 (2013).

24. Pimentel, E. C. G., Edel, C., Emmerling, R. & Götz, K.-U. How imputation errors bias genomic predictions. J. Dairy Sci. 98, 4131–4138 (2015).

25. Chen, S.-F. et al. Genotype imputation and variability in polygenic risk score estimation. Genome Med. 12, 100 (2020).

26. Khera, A. V. et al. Genome-wide polygenic scores for common diseases identify individuals with risk equivalent to monogenic mutations. Nat. Genet. 50, 1219–1224 (2018).

27. Ni, G. et al. A comprehensive evaluation of polygenic score methods across cohorts in psychiatric disorders. Genetic and Genomic Medicine (2020) doi:10.1101/2020.09.10.20192310.

28. Lee, J. J. et al. Gene discovery and polygenic prediction from a genome-wide association study of educational attainment in 1.1 million individuals. Nat. Genet. 50, 1112–1121 (2018).

29. Yengo, L. et al. Meta-analysis of genome-wide association studies for height and body mass index in ∼700000 individuals of European ancestry. Hum. Mol. Genet. 27, 3641–3649 (2018).

30. Chervova, O. et al. The Personal Genome Project-UK, an open access resource of human multi-omics data. Sci Data 6, 257 (2019).

31. Nørgaard-Pedersen, B. & Hougaard, D. M. Storage policies and use of the Danish Newborn Screening Biobank. J. Inherit. Metab. Dis. 30, 530–536 (2007).

32. Munk-Jørgensen, P. & Mortensen, P. B. The Danish Psychiatric Central Register. Dan. Med. Bull. 44, 82–84 (1997).

33. Mors, O., Perto, G. P. & Mortensen, P. B. The Danish Psychiatric Central Research Register. Scand. J. Public Health 39, 54–57 (2011).

34. Pedersen, C. B. The Danish Civil Registration System. Scand. J. Public Health 39, 22–25 (2011).

35. Schmidt, M., Pedersen, L. & Sørensen, H. T. The Danish Civil Registration System as a tool in epidemiology. Eur. J. Epidemiol. 29, 541–549 (2014).

36. Deelen, P. et al. Genotype harmonizer: automatic strand alignment and format conversion for genotype data integration. BMC Res. Notes 7, 901 (2014).

37. Purcell, S. et al. PLINK: a tool set for whole-genome association and population-based linkage analyses. Am. J. Hum. Genet. 81, 559–575 (2007).

38. Li, H. A statistical framework for SNP calling, mutation discovery, association mapping and population genetical parameter estimation from sequencing data. Bioinformatics 27, 2987–2993 (2011).

39. McCarthy, S. et al. A reference panel of 64,976 haplotypes for genotype imputation. Nat. Genet. 48, 1279–1283 (2016).

40. Choi, Y., Chan, A. P., Kirkness, E., Telenti, A. & Schork, N. J. Comparison of phasing strategies for whole human genomes. PLoS Genet. 14, e1007308 (2018).

41. Yang, J., Lee, S. H., Goddard, M. E. & Visscher, P. M. GCTA: a tool for genome-wide complex trait analysis. Am. J. Hum. Genet. 88, 76–82 (2011).

42. Chang, C. C. et al. Second-generation PLINK: rising to the challenge of larger and richer datasets. Gigascience 4, 7 (2015).

43. Price, A. L. et al. Principal components analysis corrects for stratification in genome-wide association studies. Nat. Genet. 38, 904–909 (2006).

44. Manichaikul, A. et al. Robust relationship inference in genome-wide association studies. Bioinformatics 26, 2867–2873 (2010).

45. Martin, A. R. et al. Clinical use of current polygenic risk scores may exacerbate health disparities. Nat. Genet. 51, 584–591 (2019).

46. Martin, A. R. et al. Human Demographic History Impacts Genetic Risk Prediction across Diverse Populations. Am. J. Hum. Genet. 100, 635–649 (2017).

47. >Muir, P. et al. The real cost of sequencing: scaling computation to keep pace with data generation. Genome Biol. 17, 53 (2016).

48. Das, S. et al. Next-generation genotype imputation service and methods. Nat. Genet. 48, 1284– 1287 (2016).

49. Kowalski, M. H. et al. Use of >100,000 NHLBI Trans-Omics for Precision Medicine (TOPMed) Consortium whole genome sequences improves imputation quality and detection of rare variant associations in admixed African and Hispanic/Latino populations. PLoS Genet. 15, e1008500 (2019).

## References

1. 1000 Genomes Project Consortium et al. A global reference for human genetic variation. Nature 526, 68–74 (2015).

2. Price, A. L. et al. Long-range LD can confound genome scans in admixed populations. American journal of human genetics vol. 83 132–5; author reply 135–9 (2008).

3. Purcell, S. et al. PLINK: a tool set for whole-genome association and population-based linkage analyses. Am. J. Hum. Genet. 81, 559–575 (2007).

4. Price, A. L. et al. Principal components analysis corrects for stratification in genome-wide association studies. Nat. Genet. 38, 904–909 (2006).

5. Deelen, P. et al. Genotype harmonizer: automatic strand alignment and format conversion for genotype data integration. BMC Res. Notes 7, 901 (2014).

6. Satterstrom, F. K. et al. Large-Scale Exome Sequencing Study Implicates Both Developmental and Functional Changes in the Neurobiology of Autism. Cell 180, 568–584.e23 (2020).

7. Danecek, P., McCarthy, S., Li, H. & Others. bcftools—utilities for variant calling and manipulating vcfs and bcfs. (2015).

8. Manichaikul, A. et al. Robust relationship inference in genome-wide association studies. Bioinformatics 26, 2867–2873 (2010).

9. Pedersen, C. B. The Danish Civil Registration System. Scand. J. Public Health 39, 22–25 (2011).

10. Chervova, O. et al. The Personal Genome Project-UK, an open access resource of human multiomics data. Sci Data 6, 257 (2019).

11. Pritchard, J. K. & Przeworski, M. Linkage disequilibrium in humans: models and data. Am. J. Hum. Genet. 69, 1–14 (2001).

12. Choi, S. W., Mak, T. S. & O’Reilly, P. F. Tutorial: a guide to performing polygenic risk score analyses. Nat. Protoc. 15, (2020).

